# Transcriptomic and phenomic investigations reveal elements in biofilm repression and formation in the cyanobacterium *Synechococcus elongatus* PCC 7942

**DOI:** 10.1101/2022.01.27.477154

**Authors:** Ryan Simkovsky, Rami Parnasa, Jingtong Wang, Elad Nagar, Eli Zecharia, Shiran Suban, Yevgeni Yegorov, Boris Veltman, Eleonora Sendersky, Rakefet Schwarz, Susan S. Golden

**Affiliations:** Division of Biological Sciences, University of California, San Diego, La Jolla, CA 92093, USA; The Mina and Everard Goodman Faculty of Life Sciences, Bar-Ilan University, Ramat-Gan, Israel, 5290002; Center for Circadian Biology, University of California, San Diego, La Jolla, CA 92093, USA

**Author notes:** To whom correspondence should be addressed regarding RNA-Seq and biofilm assays. To whom correspondence should be addressed regarding RB-TnSeq. Denotes equal contribution to this work (Ryan Simkovsky, Rami Parnasa).

**Keywords:** Biofilm, cyanobacteria, transposon library, RNA-Seq, RB-TnSeq

## Abstract

Biofilm formation by photosynthetic organisms is a complex behavior that serves multiple functions in the environment. Biofilm formation in the unicellular cyanobacterium *Synechococcus elongatus* PCC 7942 is regulated in part by a set of small secreted proteins that promotes biofilm formation and a self-suppression mechanism that prevents their expression. Little is known about the regulatory and structural components of the biofilms in PCC 7942, or response to the suppressor signal(s). We performed transcriptomics (RNA-Seq) and phenomics (RB-TnSeq) screens that identified four genes involved in biofilm formation and regulation, more than 25 additional candidates that may impact biofilm formation, and revealed the transcriptomic adaptation to the biofilm state. In so doing, we compared the effectiveness of these two approaches for gene discovery.

## Introduction

Biofilm formation, the adhesion of bacteria to a surface typically involving the production of an extracellular polymeric substance (EPS), is a lifestyle that allows microorganisms to survive in the face of stresses and threats from their environment, including nutrient depletion^1,2^, predation^3,4^, and dessication^5^. Biofilms can be detrimental to human health and industrial activities, by enabling persistence and antibiotic resistance in patients and on medical surfaces^6^ or by causing biofouling as on ship surfaces^7^ or in corrosion of industrial piping^8^. Studies aimed at disrupting deleterious biofilms have led to a wealth of knowledge on the molecular mechanisms regulating and forming biofilms in heterotrophic or pathogenic bacteria, including the requirement for pili and adhesins in finding and attaching to surfaces, intracellular cyclic di-GMP secondary messenger systems for controlling activation of biofilm formation, and production and secretion of exopolymeric matrix materials that can include exopolysaccharides, amyloidgenic proteins, and extracellular DNA^6^.

Photosynthetic cyanobacteria, which are among the planet’s primary producers of oxygen and fixed carbon, can also form biofilms, often in microbial mats where a combination of gas, nutrient, signaling, and light gradients determine the composition, density, and organization of the diverse community of organisms in the assemblage^9^. Cyanobacteria are becoming a prominent agricultural crop for sustainable conversion of solar energy and atmospheric carbon into replacements for petroleum products, including biofuels, industrial chemicals, nutraceuticals, and plastics^10–12^. In this context, biofilm formation can be both disadvantageous, due to biofouling of growth vessels and piping, or beneficial for protection against predation, efficiency in harvesting the crop, or minimizing resources for growth and production in a biofilm as opposed to in a planktonic culture^13,14^. In contrast to health applications that seek to prevent biofilm formation, the goal for agricultural scale industrial production of cyanobacterial biomass is to be able to reliably activate or deactivate biofilming to minimize costs and damage to equipment while maximizing productivity.

Only in recent years have studies on the molecular mechanisms of biofilm formation extended to cyanobacteria, where light limitation through self-shading is an inherent consequence of biofilm structure^13^. For many cyanobacteria, biofilm formation and regulation appears to have a number of features in common with heterotrophic bacteria: type IV pili and S-layer proteins are required for film formation in *Synechocystis* sp. PCC 6803^15,16^, cyclic-di-GMP promotes aggregation in *Thermosynechococcus vulcanus*^17^ and biofilm formation in PCC 6803^18^, a two-component regulatory system comprising a response regulator and a split histidine kinase controls cell aggregation in PCC 6803^19^, and extracellular polysaccharides, such as cellulose in *T. vulcanus*, contribute to the extracellular matrix and cellular aggregation^20^.

In contrast, our studies have demonstrated that biofilm formation in *Synechococcus elongatus* PCC 7942, which is by default constitutively suppressed in the laboratory^21^, does not require pili^22^. In fact, most of the mutations discovered to date that relieve suppression and enable biofilm formation (Supplemental Table S1) block formation of the Type IV pilus or its assembly apparatus, including the pilin-encoding *pilA1* and *pilA2* genes; the pilus assembly and Type II secretion proteins encoded by *pilB* (formerly referred to as *t2sE*), *pilC*, and *pilN*; and two proteins encoded by the genes *hfq* and *ebsA* that form a tripartite complex with PilB^21–23^. For the majority of these mutants, a biofilm containing 90 – 95% of the cell population forms 3 – 4 days following inoculation in bubbling tubes. Electron microscopy has confirmed that the biofilm cells lack pili. Furthermore, they have markedly diminished or altered exoproteomes, suggesting that a single Type IV/II pilus and secretion system fills these dual roles in *S. elongatus*^21–23^. Removal of this complex is pleiotropic – impeding other ecologically-relevant behaviors such as natural competence and phototaxis-directed twitching motility^24,25^.

Wild-type *S. elongatus* PCC 7942 (WT) does not produce biofilms under laboratory conditions due to the production and secretion of a self-suppressor^21^. WT growth medium (conditioned medium, CM) contains accumulated suppressor that prevents biofilm formation by the *pilB* mutant, *PilB*::Tn*5* (formerly named T2SEΩ), but CM from *PilB*::Tn*5* does not, indicating that the mutant still responds to the suppressor it does not secrete^21^. The derepression of a set of four small secreted proteins (EbfG1-4), each characterized by a secretion motif shared with microcins, enables biofilm formation in *PilB*::Tn*5*^21,26,27^. The processing and secretion of these small proteins requires the cysteine peptidase gene *pteB*, which is cistronic with the *ebfG* genes, and the putative processing peptidase gene *ebfE*, whose inactivation significantly alters the exoproteome differently than changes observed with inactivation of *pilB*^27^.

The robust formation of biofilms by the recent wild isolate *S. elongatus* UTEX 3055 demonstrates that biofilm formation of PCC 7942 mutants uncovers a process that is reversible in the natural environment^25^. Decades of laboratory culturing techniques, such as decanting, that subtly select for planktonic cells^28^ has likely domesticated the PCC 7942 WT strain into a biofilm-suppressed state through genetic drift or epigenetic changes, which now requires biofilm-inducing mutations to activate the biofilm program.

The known gene products involved in suppressing or enabling biofilm formation in *S. elongatus* PCC 7942 (Supplemental Table S1) represent only pieces of the structural and regulatory system of biofilm formation, which must include the circuitry to synthesize, detect, and respond to the secreted repressor, as well as the repressed components that contribute to biofilm formation once self-suppression is alleviated. To obtain a more comprehensive understanding of genes involved in biofilm formation we leveraged the natural competence^29^ of *S. elongatus* and genetic tools that include a library of transposon mutagenesis vectors for the rapid individual knockout of nearly any non-essential gene in the genome^30,31^ and a randomly barcoded transposon sequencing (RB-TnSeq) library for quantitatively assessing fitness values and directly linking genotypes with phenotypes^32,33^. Analogous TnSeq libraries in other organisms have been used to study complex behaviors, as in biofilm formation in *Bacillus cereus*^34,35^ or in community interactions of bacteria and fungi on the surface of maturing cheese^36,37^. In addition to phenomics using RB-TnSeq, which identifies genes whose loss either enhances or reduces a cell’s fitness in a biofilm, we employed transcriptomics using RNA-Seq, which identifies genes whose transcription differs between planktonic and filmed cells. These two data sets suggest that RB-TnSeq is more powerful for identifying genes that contribute to a phenotype, whereas transcriptomics describes the underlying biology of a phenotypic state.

## Materials and Methods

### Culture conditions, plasmids, and strains

For standard growth of *S. elongatus* without additional CO_2_, cultures were grown at 30 °C in BG-11 medium^38^. For biofilm assays performed in test tubes, where cultures were bubbled with 3% CO_2_, or in 96-well plates, cultures were inoculated into BG-11 media with HEPES and freshly prepared ferric ammonium citrate and citric acid components as previously described^39^. Preparation and collection of conditioned media from WT cultures grown in BG-11 was performed as described previously^21,26^.

All vectors used in this study are listed in Supplemental Table S10. Except for the RB-TnSeq experiments, where strains were derived from the Golden lab culture collection, all *S. elongatus* strains were derived from those in the Schwarz lab culture collection, which lack the small plasmid, pANS. Generation of novel insertional knockout *S. elongatus* strains using Unigene Set (UGS)^30,31^ or the constructed vector EZ27 were performed according to standard transformation methods that take advantage of *S. elongatus*’ natural competence and ability to perform double homologous recombination^40^. In cases in which other mutations were combined with a *pilB* disruption, *pilB* inactivation was introduced after knockout of the other gene of interest due to impaired competence in *PilB*::Tn*5*^21^. Double recombination and segregation of mutants were confirmed via PCR using primers listed in Table S10.

An inactivation vector for gene Synpcc7942_0774 was constructed through blunt-end cloning of a PCR-amplified gene fragment, generated using the K273 and K274 primers listed in Table S10, into the pJet1.2 vector (Thermo Fisher Scientific). A spectinomycin-resistance cassette was inserted at the XcmI site in the gene fragment, yielding the insertional knockout construct (pEZ27) that was used for transformation and homologous recombination into WT.

### Biofilm formation assays

Unless noted otherwise, growth of *S. elongatus* PCC 7942 and all derived strains for the purpose of generating biofilms were performed in bubbling tube cultures containing 25 ml of BG-11 with HEPES after inoculation, using fresh starter cultures derived from solid media cultures, at an OD_750_ of 0.5 at 30 °C and under low light (5 – 30 µmol photons m^−2^ sec^−1^), as described previously^39^. Biofilm assays under static conditions were grown in 96-well plates, as previously described^27^, or in 5 ml culture volumes in sterile 25 ml Pyrex glass flasks (# 4980) plugged with cotton balls that were incubated under low light (5 – 30 μmol photons m^−2^ sec^−1^) without shaking for 6 days prior to biofilm quantitation. Quantification of the biofilm formation as a percentage of chlorophyll in suspension or using ethanol-extracted crystal violet staining was performed as described previously^27,39^.

### RNA-Seq Experiment

WT and *PilB*::Tn*5* strains were grown for biofilm formation in triplicate sets of bubbling tubes as described previously^26^. Samples were taken for RNA extraction from the liquid supernatants one and four days after inoculation (Days 1 and 4). For the *PilB*::Tn*5* culture samples on Day 4, samples were first taken from the supernatant fraction, after which the liquid was removed by decanting, the tubes were gently washed with deionized water, and the remaining biofilm left behind was scraped to obtain biofilm samples. Total RNA was extracted as previously described by Tu et al.^41^, with the following modifications: phenol was saturated with water; cell extraction was performed by vortexing for 1 min, incubating at 65 °C for 10 min with a brief vortex after 5 min, and incubating on ice for 3 min. The beads and unbroken cells were pelleted by centrifugation (Eppendorf centrifuge, maximal speed, 10 min), and the aqueous phase was subjected to one phenol, two to three phenol-chloroform (1:1), and one chloroform extraction, where chloroform was pre-equilibrated with isoamyl alcohol at 24:1 ratio. Nucleic acids were precipitated by adding 1/3 volume of 8 M LiCl and 2 volumes of ethanol followed by gentle mixing, incubation for 40 min at −80°C or for 2 h at −20°C, and centrifugation as described. The pellet was washed with 70% ethanol and resuspended with HPLC-grade water. 40 µg of total RNA was treated with 4 U of TURBO(tm) DNase (Ambion, Catalog #: AM2238) at room temperature for 45 min followed by a boost with 4 U and an additional 45-min incubation. The DNase was removed by phenol-chloroform and chloroform extractions and the RNA was precipitated as described above. RNA sample pellets (21), representing triplicates of the seven sampling groups described above and resulting from the ethanol precipitation, were washed with ice cold 70% ethanol. The resulting air-dried pellets were resuspended in RNase-free water and 3 µg of DNA from each sample was depleted of ribosomal RNA using the Bacteria Ribo-Zero rRNA Removal Kit (Epicentre) and purified using the RNeasy miniprep kit (Qiagen), both performed according to the manufacturer’s instructions. Recovery rates for each sample fell in the range of 3.6 – 9.2% and rRNA depletion and RNA quality was confirmed for a subset of samples using 1% agarose gel electrophoresis. Purified ribosomal RNA depleted samples were submitted to the Scripps Research (La Jolla, CA) sequencing core for strand-specific library preparation and sequencing on an Illumina 1×100 TruSeq system.

### RNA-Seq Data Analysis

FASTQ files were trimmed using trimmomatic^42^ prior to analysis of the seven sampling groups with two separate RNA-Seq analysis pipelines: 1) Rockhopper 2^43^, an all-in-one analysis tool specifically designed for analysis of bacterial transcriptomics data, using default parameters and 2) A custom scripted pipeline written in shell commands and in R^44^ using Bowtie2^45^ for alignment to the genome, SAMtools^46^ and BEDTools^47^ for file format conversion and generation of coverage files for visualization in IGV^48^, htseq-count for generating strand-specific gene counts^49^, the DeSeq2^50^ R package for differential gene expression analysis using an alpha value of 0.05 and either a likelihood ratio test for ANOVA analysis of multiple sample groups simultaneously using or the Wald significance test for pairwise sample group comparisons, the ashr^51^ R package for determining effect size shrinkage estimates, and tidyverse^52^ R packages for further data organization and visualization. For both pipelines, significantly differentially expressed genes were determined for each of the 21 possible pairwise comparisons of the 7 sample groups using a false detection rate (FDR; q-value from Rockhopper2 and FDR-adjusted p-values from DESeq2) threshold of ≤ 0.05 to determine significance and a |log_2_(Fold Change)| > 1 for differentiating highly changed genes, labeled “ Up” or “ Down”, from genes with minor expression changes, labeled “ Neutral Up” or “ Neutral Down”. All alignment statistics, gene counts, fold change values, FDR values, and DEG assignment data for the Rockhopper2 analysis are provided in Supplementary File 1, while the same data for the DESeq2-based analysis are provided in Supplementary File 2. To consolidate the lists of DEGs for each pairwise comparison from each analysis pipeline, a union of the data sets was performed so that the labels “ Up” or “ Down” were applied to any gene in which the gene was so labeled in at least one of the data sets, while “ Neutral Up” or “ Neutral Down” was applied to any remaining gene that was otherwise determined as significantly changed using the FDR threshold in either of the analyses. Comparisons of pairwise comparisons were then performed by manually clustering genes into groups of interest with similar patterns of DEG categorization across relevant pairwise experiments. Groupings that consider the “ Neutral” genes (labeled “ Full”) are provided in Supplemental File 3, as well as the groupings discussed herein that do not consider the “ Neutral” genes for categorization purposes (labeled “ Reduced”). These latter groups of interest served as the basis for the enrichment analysis detailed below. All data regarding the pipeline analyses consolidation, DEG categorization, comparisons of pairwise comparisons, and enrichment analysis are provided in Supplemental File 3.

### RB-TnSeq Experiment

For the first experiment, an aliquot of the *S. elongatus* RB-TnSeq version 1.0 library^32^ was revived and grown in three flasks of 100-ml BG-11, as described previously^24,53^. The three flasks were pooled to generate a culture with OD_750_ of 0.4. An aliquot of this pooled culture was pelleted by centrifugation and frozen at −80°C to serve as a T0 sample. A culture (93 ml) of the pooled library was centrifuged at 5,000 rpm at room temperature for 15 min. The pelleted cells were resuspended in 75 ml of BG-11 with HEPES media to an OD_750_ of 0.5, split equally into triplicate bubbling tubes, and allowed to form biofilms for 21 days at 30 °C and under low fluorescent light (5 μmol photons m^−2^ sec^−1^) while bubbled with hydrated 3% CO_2_.

For the second experiment, an aliquot of the *S. elongatus* RB-TnSeq version 2.0 library was revived, grown, and sampled for a T0 sample as performed for the version 1.0 library. Aliquots of the library were centrifuged to pellet cells and resuspended in either fresh or WT conditioned BG-11 media, both with HEPES, to an OD_750_ of 0.5. Resuspensions were then split into triplicate bubbling tubes containing 25 ml of culture each or into sterile 25 ml Pyrex glass flasks (# 4980), plugged with cotton balls, containing 5 ml of culture each. Bubbling tube cultures for this experiment were placed under the same conditions as for the first experiment for 15 days to allow biofilms to form. Flask cultures were incubated under low fluorescent light (5 μmol photons m^−2^ sec^−1^) without shaking at room temperature for 15 days.

After the appropriate incubation periods, biofilm formation was confirmed visually. Planktonic cell samples were collected by decanting the supernatants of the cultures into sterile conical tubes. Each vessel was then gently washed once with water, which was decanted to a sterile conical tube to investigate settler cells. The remaining biofilms were resuspended in a water wash through physical scraping of the vessel and collected into a sterile conical tube. All samples were then centrifuged to generate cell pellets, which were frozen at −80°C prior to genomic DNA extraction^40^, barcode PCR amplification using Q5 DNA polymerase (NEB) and Illumina adaptor-encoded primers that include unique 6-bp TruSeq indexes in the forward primer, PCR purification of barcode amplicons with the DNA Clean and Concentrator kit (Zymo Research), and Illumina HiSeq single-end sequencing at LBNL, all as detailed in previously reported RB-TnSeq protocols^54^.

### RB-TnSeq Data Analysis

Barcode sequences were extracted and counted from FASTQ files using PERL scripts previously reported by Wetmore, et al.^54^ using an updated genome annotation that includes the known biofilm enhancing *ebfG* genes and other ORFs that are not annotated in publicly available genome files. Strain fitness, gene fitness, and T-value *t*-like statistics were calculated and a T-value threshold for significance of ≥ 4 was applied as in Wetmore, et al.^54^. Significant genes with an absolute fitness score > 1 were labeled “ Up” or “ Down”, while those with absolute fitness scores ≤ 1 were labeled “ Neutral Up” or “ Neutral Down”.

### Gene Enrichment Analysis

Gene categorical information, including known biofilm genes from previous *S. elongatus* PCC 7942 biofilm publications^22,23,26,27,55^, pili genes described by Taton, et al.^24^, essentiality and conservation data from Rubin, et al.^32^, functional categories from the JGI IGM^56^ and CyanoBase^57^ databases, and signal peptide and secretion predictions^58–63^, were amassed for all genes in the *S. elongatus* PCC 7942 genome from appropriate sources and are provided in Supplemental Tables S1-3. Significant enrichments of genes in each interest group identified for RNA-Seq and RB-TnSeq data comparison sets for each of the informational categories were determined using custom R scripts to perform two-sided Fisher’s exact tests, with FDR-adjusted p-values ≤ 0.05 being designated as significant. Fold enrichment (F) was calculated as the number of genes in the interest group that are also in the category (N_gc_) divided by the number of genes expected in the group and category (E_gc_). This expected number was calculated by multiplying the number of total genes in the interest group (N_g_) by the frequency of all genes in the genome that are found in the category (f_c_), which was determined as the number of genes in the category (N_c_) divided by the number of genes in the genome (N).

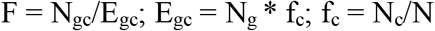

## Results and Discussion

### RNA-Seq characterization of WT and biofilm-forming strains

Our first approach aimed to identify genes that are differentially regulated between planktonic and biofilm states. To compare the transcript profiles of cells destined to form or actively forming biofilms with those of planktonic cells, as well as the impact that conditioned (biofilm-suppressing) medium has upon these processes, total RNA was purified from WT or *PilB*::Tn*5* cultures inoculated in either fresh or WT-derived conditioned medium (CM) and sampled 1 and 4 days following inoculation. Only the 4-day *PilB*::Tn*5* cultures grown in fresh BG-11 produced biofilms, so transcript profiles were processed from both planktonic and biofilmed fractions from these cultures (Fig. 1A). Twenty-one samples representing triplicate experiments of seven experimental conditions were depleted of ribosomal RNA and sequenced using an Illumina platform, yielding 6 – 8 million reads per sample. Data were aligned to the *S. elongatus* PCC 7942 genome and analyzed for differentially expressed genes (DEG) using two different software pipelines: 1) Rockhopper 2, a prokaryotic-specific RNA-seq analysis platform^43^ and 2) a pipeline of published RNA-Seq softwares including Bowtie2^45^ for alignment, htseq-count^49^ for gene counts, and DESeq2^50^ for DEG analysis, with each analysis pipeline aligning approximately 99% of trimmed reads for nearly all samples, with less than 1% of these reads mapping to ribosomal RNAs (See Supplemental Files S1 and S2 for further details). Both the WT and *PilB*::Tn*5* strains used in these experiments lack the smaller plasmid, pANS, and thus its potential impact on biofilm formation was not assessed.

**Figure 1:**
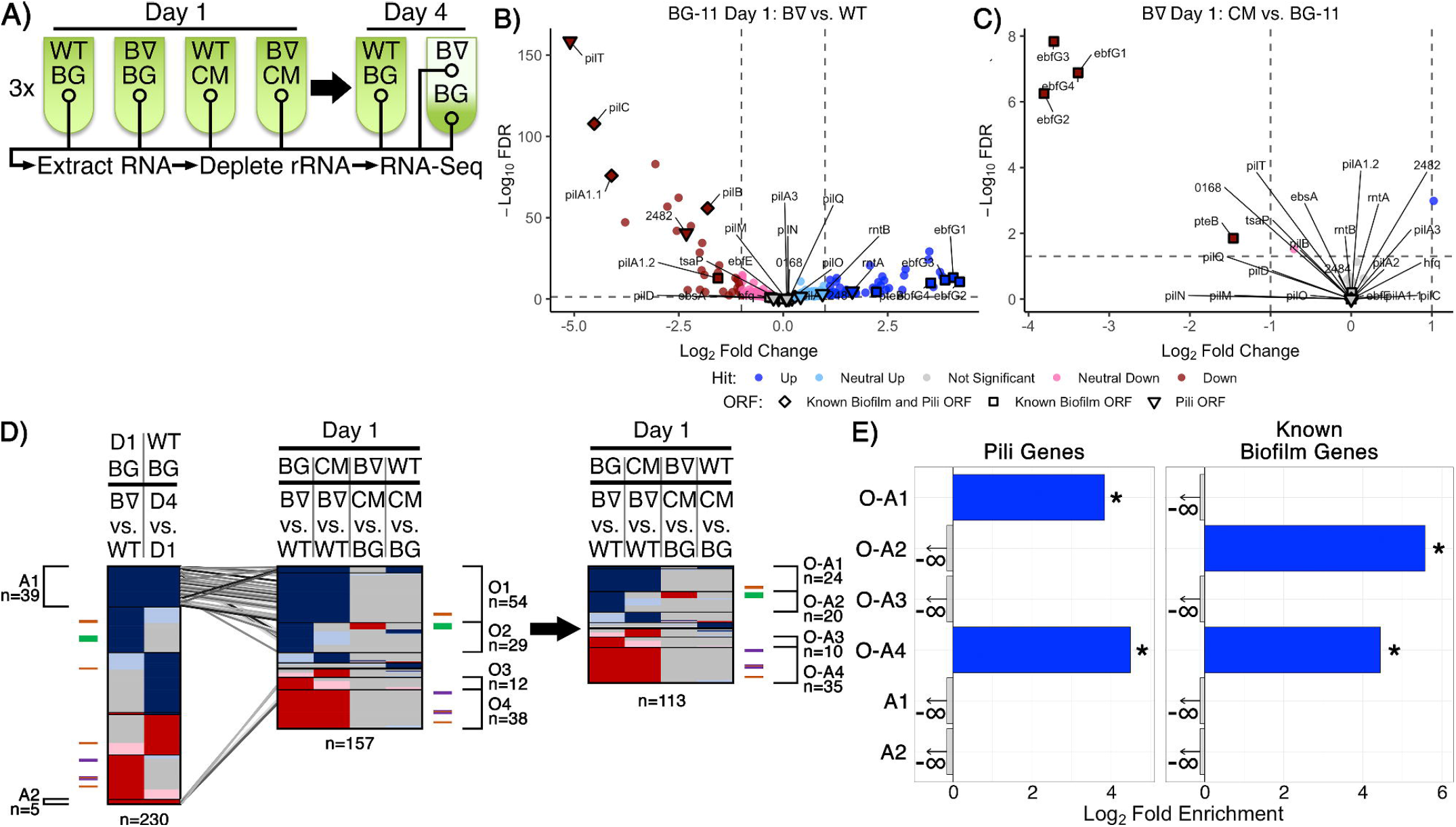
RNA-Seq of WT and *PilB*::Tn*5* and Day 1 DEG analysis. A) Diagram of the time course for triplicate samples processed for Wild Type (WT) and *PilB*::Tn*5* (B∇) when placed under biofilm forming conditions in either fresh BG-11 (BG) or WT-derived conditioned media (CM). Samples (circles) include planktonic samples from all test tubes and a sample of the biofilm formed by *PilB*::Tn*5* on Day 4 in BG-11. B & C) Volcano plots comparing the log_2_ fold change of pairwise comparisons of samples against their −log_10_ false discovery rate (FDR) as calculated by DESeq2. Data points are colored based on their DEG designation (“ Hit”). Labeled shapes indicate known biofilm and pili ORFs. Dashed lines indicate the thresholds used for hit designation. Note that each plot is scaled individually. D) Comparisons of sDEG designations (colored as in B & C) of ORFs (y-axis) from different relevant pairwise comparisons, as labeled above. The locations in the graph of known pili and biofilm ORFs are indicated by the colored ticks next to each graph (purple = biofilm and pili ORF, orange = pili ORF, green = biofilm ORF) and clusters of genes with common sDEG patterns across the x-axis of interest are designated with brackets and labeled with their identifiers and the number of sDEGs in the group. The number of ORFs considered in each graph is provided below the graph. On the left, the Day 1 BG-11 *PilB*::Tn*5* versus WT pairwise comparison is compared against the WT BG-11 Day 4 vs. Day 1 comparison to identify ORFs related to the stress or density state of the culture. Those ORFs that are similarly up- or down-regulated in these two pairwise comparisons (groups A1 and A2) are connected to their location in the middle graph comparing all relevant Day 1 pairwise comparisons by the diagonal lines. These ORFs were then removed from the middle graph to generate the graph on the right. E) Enrichment analyses of pili genes and known biofilm genes in the O-A and A groups of interest labeled in D. Significant enrichment (FDR ≤ 0.05) is designated by solid bars and stars. Groups that do not have any of the genes analyzed are designated with a negative infinity and arrow, as appropriate for the log_2_ scale of the x-axis.

ANOVA analysis using DESeq across all samples demonstrated that over 65% (1776 out of 2729) of the ORFs encoded by the genome, including all known biofilm-related (Table S1) and pili (Table S2) genes except for *tsaP*, are differentially expressed between at least two experimental conditions (File S2 Sheets 4 & 8). Additionally, Rockhopper predicted 118 non-coding RNAs (ncRNAs), of which 42 are antisense to annotated ORFs, 55 overlap with or are enclosed by previously reported potential ncRNAs^64,65^, and one overlaps an essential intergenic region^32^ (Table S4, File S1 Sheet 9). All but three of these predicted RNAs were differentially expressed between at least two experimental conditions. Although RNA-binding by cyanobacterial Hfq homologs so far has not been demonstrated^66^, we hypothesize that these novel differentially expressed RNAs may represent binding targets.

Both Rockhopper and DESeq analyses were performed for all twenty-one pairwise comparisons of the seven experimental groups (Files S1 Sheet 6 and S2 Sheet 5). The union of the lists of DEGs from the two analyses generated a final list of DEGs for all pairwise comparisons (File S3 Sheets 1 & 2); the intersection of the lists proved too stringent and excluded known biofilm forming genes that the union did not.

Ten pairwise comparisons (Figs. 1, 2, & S2; File S3 Sheet 4) are relevant to the experimental variables tested: the *pilB* genetic alteration, biofilm state, time, and media conditions. While recognizing that more modest changes in expression may be biologically relevant, focusing on genes with fold changes > 2, designated as strong DEGs (sDEGs), limited the total set considered among the ten pairwise comparisons nearly four-fold from 1,867 DEGs to 508 sDEGs and resulted in a more limited range of sDEGS per pairwise comparison of 10 and 18 for those involving CM to 344 for *pilB*::Tn*5* biofilm formation over time (Figs. S1A,C; File S3 Sheets 2 & 4), in contrast to 15, 67, and 1332 DEGs, respectively.

**Figure 2:**
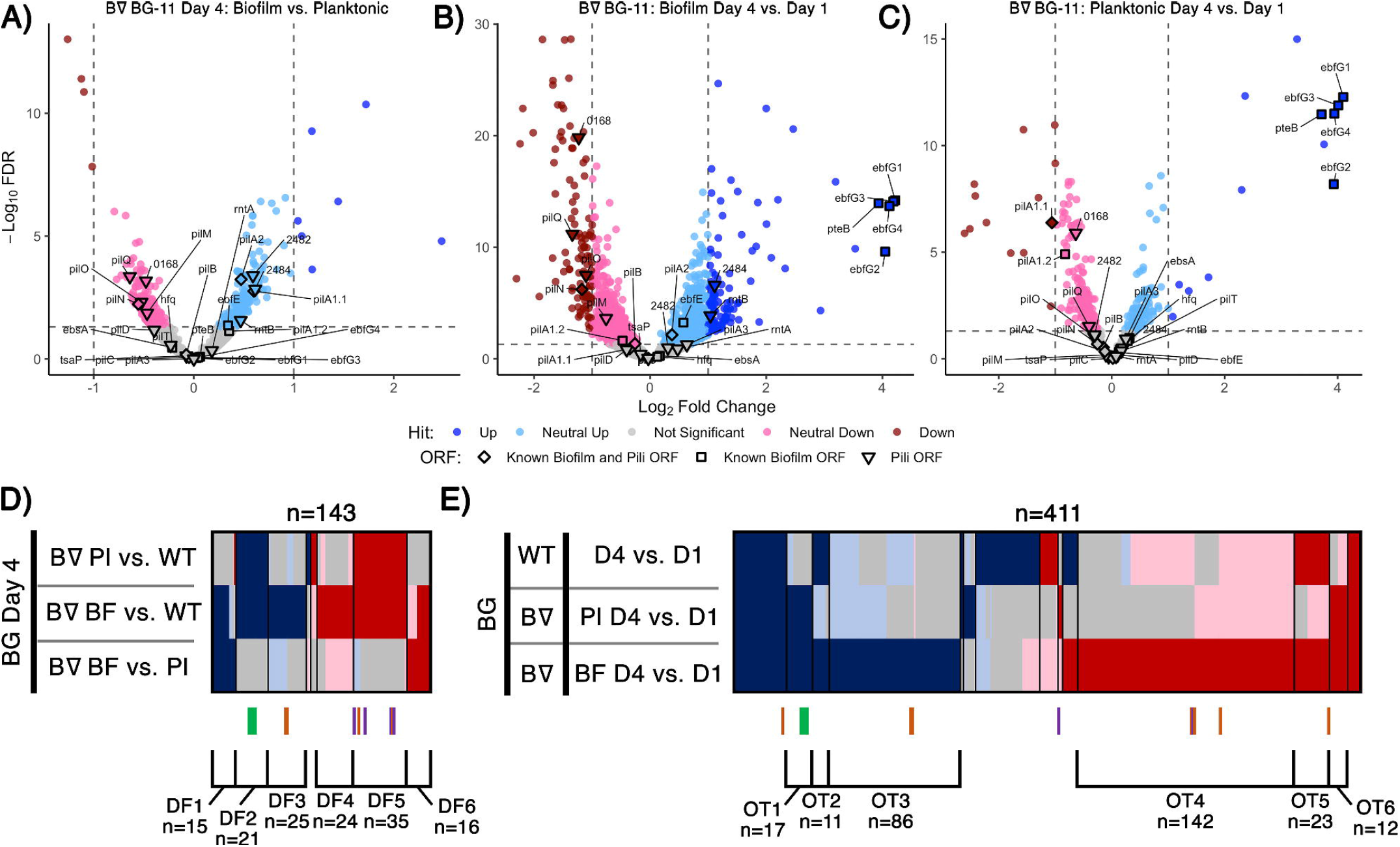
Day 4 and time course RNA-Seq analysis. Volcano plots, presented as described for Figs. 1B,C, of A) Day 4 pairwise comparison of *PilB*::Tn*5* biofilms versus planktonic cells, B) *PilB*::Tn*5* biofilms on Day 4 versus *PilB*::Tn*5* on Day 1, and C) *PilB*::Tn*5* planktonic cells on Day 4 versus *PilB*::Tn*5* on Day 1. Comparison of sDEGs associated with D) Day 4 and E) over time are presented as described for Fig. 1D.

Tracking the 29 known biofilm and pili genes (Table S1 and S2, File S3 Sheet 6) enabled evaluation of the biological significance of these results. All except 7 (*tsaP, pilD, hfq, ebfE, ebsA, pilA2* and the second *pilT* gene) were identified as sDEGs (Figs. 1B,C,D & S2, File S3 Sheet 6). While it is reasonable that *tsaP, pilD*, and the second *pilT* genes do not have notable impacts on biofilm formation due to degeneracy or their encoded proteins’ roles as accessories to the pilus, the genes of the Hfq-EbsA-PilB tripartite complex^23^ or the EbfE-PteB^26^ maturation-secretion system play important roles that likely act at post-transcriptional levels^23,55^. These results demonstrate the limitations of using changes in expression level as a proxy for contribution to a particular phenotype.

### Transcriptome comparisons of biofilm and planktonic cells reflect the biofilm-proficient *pilB*::Tn*5* genotype

We initially hypothesized that the Day 4 comparisons between biofilm and planktonic samples would identify novel genes required to regulate and create the biofilm. However, the sDEG patterns instead point to transcriptional consequences of the *pilB*::Tn*5* mutation independent of its biofilm state or to the change in environment associated with existing in the biofilm (Figs. 2, S2, & S3). None of the known biofilm and pili genes differed sufficiently in expression between *pilB*::Tn*5* biofilms and planktonic cells to be considered as sDEGs (Fig. 2A, File S3 Sheet 6), but the majority of them were either up-regulated (e.g *ebfG* genes, named group DF2) or down-regulated (e.g. *pilB* operon genes, group DF5) when comparing either of the *PilB*::Tn*5* samples against WT (Fig. 2D). The sets of similarly up- or down-regulated sDEGs in DF2 and DF5 are enriched for genes related to defense mechanism, motility, and proteins with predicted secretion signals (Fig. S6B,E, File S3 Sheets 10 & 18), consistent with known alterations in the exoproteome of *PilB*::Tn*5*^22,23,27^.

Sets of 15 up-regulated and 16 down-regulated sDEGs correlate with the biofilm state (groups DF1 and DF6, respectively, in Fig. 2D, File S3 Sheet 10). The set of down-regulated sDEGs (DF6) is enriched for Cyanobase categories of energy and inorganic ion metabolism (Figs. S5A & S6D, File S3 Sheet 18), suggesting that biofilm cells damp down core photosynthetic, ATP synthesis, nitrate, and iron nutrient metabolisms. Up-regulated sDEGs (DF1) may reflect the resource limitations and self-shaded state of the biofilm. DF1 is dominated by genes encoding hypothetical proteins (File S3 Sheets 10 & 18) and include: *sigD/rpoD3* (sigma factor), which is typically induced under high light in PCC 7942^67^ and is also associated with survival under nitrogen deficiency^68^ and resistance to oxidative stress in PCC 6803^69^; and the phycobilisome degradation protein *nblA*. Other DF1 biofilm-specific up-regulated sDEGs relate to stress responses including metal limitations, such as the high-light inducible *hliA*^70^ and genes encoding a putative ferric uptake regulator, a cyclic-AMP binding protein, and a cupredoxin.

Day 1 and Day 4 samples were compared to identify genes that might be involved in the process of biofilm formation. Genes in the *ebfG*-cluster and *pteB* are the most up-regulated sDEGs in both *PilB*::Tn*5* biofilm and planktonic samples (Figs. 2B,C) and are the only non-pili biofilm sDEGs over time. Although the distributed pattern of pili sDEGs over time does not highlight any specific cluster of candidates (Fig. 2E), the set of genes that are up-regulated similarly to the *ebfG*-cluster in both *PilB*::Tn*5* samples over time (group OT1) is enriched with 9 genes of unknown function (Fig. S7B, File S3 Sheet 12) that may play a role in biofilm formation. Biofilm *PilB*::Tn*5* cells result in a much larger number of sDEGs (148 up; 196 down) than planktonic cells (61 up; 24 down) when compared to Day 1 samples (Figs. 2B,C). Assessment of all over-time pairwise comparisons (Fig. 2E, File S3 Sheet 12) highlights 86 up (group OT3) and 142 down (group OT4) biofilm-specific sDEGs, whose overlapping composition with the sDEGs of the Day 4 comparisons and an OT4-specific enrichment for energy metabolism, motility, and unknowns (Fig. S5B, File S3 Sheet 18) reinforce that the major changes in transcript abundance over time relate to adaptation to the self-shaded, nutrient-limited biofilm condition, rather than induction of components that control or form the biofilm phenotype.

### Day 1 transcriptomics highlight regulatory and CM-responsive genes of interest

The four pairwise comparisons of all Day 1 samples identified nearly all known biofilm and pili genes among the highest magnitude up- and down-regulated sDEGs in the comparison of *PilB*::Tn*5* with WT (Fig. 1B). Whether in fresh (Fig. 1B) or conditioned media (Fig. S2), the *pilB*::Tn*5* mutation strongly decreased structural pilin gene expression, including *pilB* and downstream genes. The Day 1 data reliably capture the known genotype-phenotype associations related to biofilm regulation, as the *ebfG* genes were the most upregulated genes in *PilB*::Tn*5* compared to WT in fresh media *ebfG* genes (Figs. 1B & S2), and were strongly down-regulated (Fig. 1C) in CM. The 54 up-regulated (group O1) and 38 down-regulated (group O4) sDEGs that differ between *pilB*::Tn*5* and WT on Day 1 (Fig. 1D, File S3 Sheet 8), independent of the media, are comprised of numerous Day 4 sDEGs that are associated with responses to the environment. These include up-regulation of heat shock proteins Hsp20 and DnaK and an iron-stress chlorophyll-binding protein, and down-regulation of heat shock protein GrpE. Some of these genes are sDEGs for WT when examined over time, suggesting that a pleiotropic effect of the *pilB*::Tn*5* mutation is early establishment of a condition that WT assumes in a more advanced culture. These sDEGs suggest that *PilB*::Tn*5* statically resembles a stationary culture, as these genes are not sDEGs when this mutant is examined over time.

To filter out the sDEGs related to this apparent constitutive-stress state, we compared the sDEGs set for *PilB*::Tn*5* versus WT in BG-11 on Day 1, which reflects the genetic difference between the strains, against the sDEGs set for WT in BG-11 on Day 4 versus Day 1, which presumably reflects response of the cells to aging of the culture and diminishing resources in low light-penetration conditions (Fig. 1D, left). This comparison identified 39 (group A1) up- and 5 (group A2) down-regulated sDEGs in both pairwise comparisons, which are enriched in proteins of unknown function and Phobius-predicted secretion signals, respectively (Figs. S4A,B, File S3 Sheets 14 & 18). Group A1 includes stress-related genes, such as *hsp20* and the iron-stress chlorophyll-binding protein previously noted, as well as transcription regulators such as *sigF2*, which regulates dark-induced competence^24^ and is associated with pili formation and desiccation tolerance in other cyanobacteria^71^, and a Crp/Fnr transcriptional family regulator. These sDEGs support the premise that stress management genes associated with alterations in pili and protein maintenance that occur in a dense, low-light culture are regulated in WT, but are constitutively expressed at stress-response levels at all times in *PilB*::Tn*5*.

Based on this analysis, genes in groups A1 and A2 were removed from the Day 1 comparison and four primary groups of interest were identified as being up- or down-regulated in *PilB*::Tn*5* independent of the media (O-A1 and O-A4, respectively) or in fresh BG-11 media only (O-A2 and O-A3, respectively) (Fig. 1D, File S3 Sheet 16). While O-A1 is not further enriched for any particular functionality beyond that of pili genes (Figs. 1E & S4C, File S3 Sheet 18), this group of 24 sDEGs includes genes encoding two response regulator proteins, a histidine kinase, a glycosyltransferase, and 13 genes of unknown function that may contribute to the regulation of biofilm formation. O-A2 is enriched in known biofilm genes, predicted secreted proteins, and includes a response regulator protein, a lipoyltransferase, and 11 genes of unknown function that may participate in biofilm production along with the EbfG proteins (Figs. 1E & S4D, File S3 Sheet 16). These up-regulated clusters of sDEGs highlight potential novel genes involved in biofilm formation.

Group O-A4 is enriched in known pili and biofilm genes and genes encoding proteins of unknown function, proteins involved in motility, and predicted secreted proteins (Figs. 1E & S4F, File S3 Sheets 16 & 18); these properties are consistent with potential novel biofilm-repressing genes. Two genes of interest that encode proteins with cyclic di-GMP-binding domains and GAF domains, Synpcc7942_1148 and Synpcc7942_2096, may contribute to regulation of biofilm formation through cyclic di-GMP signaling, as occurs in *Synechocystis* PCC 6803^18^.

The pairwise comparisons pertaining to the impact of conditioned media, Day 1 in BG-11 versus CM for both *PilB*::Tn*5* and WT, resulted in small sets of 10 sDEGs each (Table S6 groups IG1 & IG2), with the most prominent impact of CM being the down-regulation of the *ebfG* operon genes in *PilB*::Tn*5* (Fig. 1C), consistent with previous RT-PCR data^26^. Most of these genes encode proteins of unknown function (File S3 Sheet 16).

Overall, the RNA-Seq experiment provides an overview of the pleiotropic phenotypes associated with the *PilB*::Tn*5* mutation and a large set of genes that likely reflect the impact of the environment of the biofilm, which is self-shaded and has limited gas and nutrient exchange, rather than the active process of biofilm formation. Extensive comparative analysis does identify sDEG clusters of interest that include candidate genes for biofilm regulation, formation, and responses to conditioned media, though the majority of these potential novel biofilm genes encode proteins of unknown function and must be investigated through other means.

### RB-TnSeq screen for loci that affect biofilm formation

A complementary approach used an RB-TnSeq assay that measures the contribution of any given gene in the genome to influence the ability of a cell to proliferate in biofilms versus planktonic growth. We screened a pooled RB-TnSeq library of approximately 150,000 individual barcoded insertions mapped to the *S. elongatus* PCC 7942 genome^32^. PCR amplification and high-throughput sequencing of the barcodes quantifies changes in abundance of individual insertion mutants in the population under varying conditions, directly associating phenotypes with genotypes as has been shown for diurnal growth^53^, c-di-AMP utilization^72^, and natural competence^24^. We grew replicate biological samples of the RB-TnSeq library under a variety of biofilm formation conditions, including in bubbling tubes and stationary flasks in fresh BG-11 or CM, in two separate experiments (Fig. 3A). Conditions were similar to those used for RNA-Seq experiments except that biofilm formation was allowed to proceed for two-weeks to ensure sufficient biomass to perform genome extraction and barcode counting. Because the majority of the mutant cells in the library should still secrete and respond to the biofilm-repressing signal that is characteristic of the WT and its CM^21^, it was not expected *a priori* that the library could form biofilms. Therefore, we grew samples of a characterized biofilm forming mutant and a 1:1 mixture of WT and the biofilm-forming mutant in parallel with the library as controls. The transposon-mutant library samples, but not the WT, generated visible biofilms, although never to the extent observed for the biofilm mutant (Fig. 3B). The 1:1 mixture of the biofilm-forming mutant and WT also produced an intermediate level of visible biofilm.

**Figure 3:**
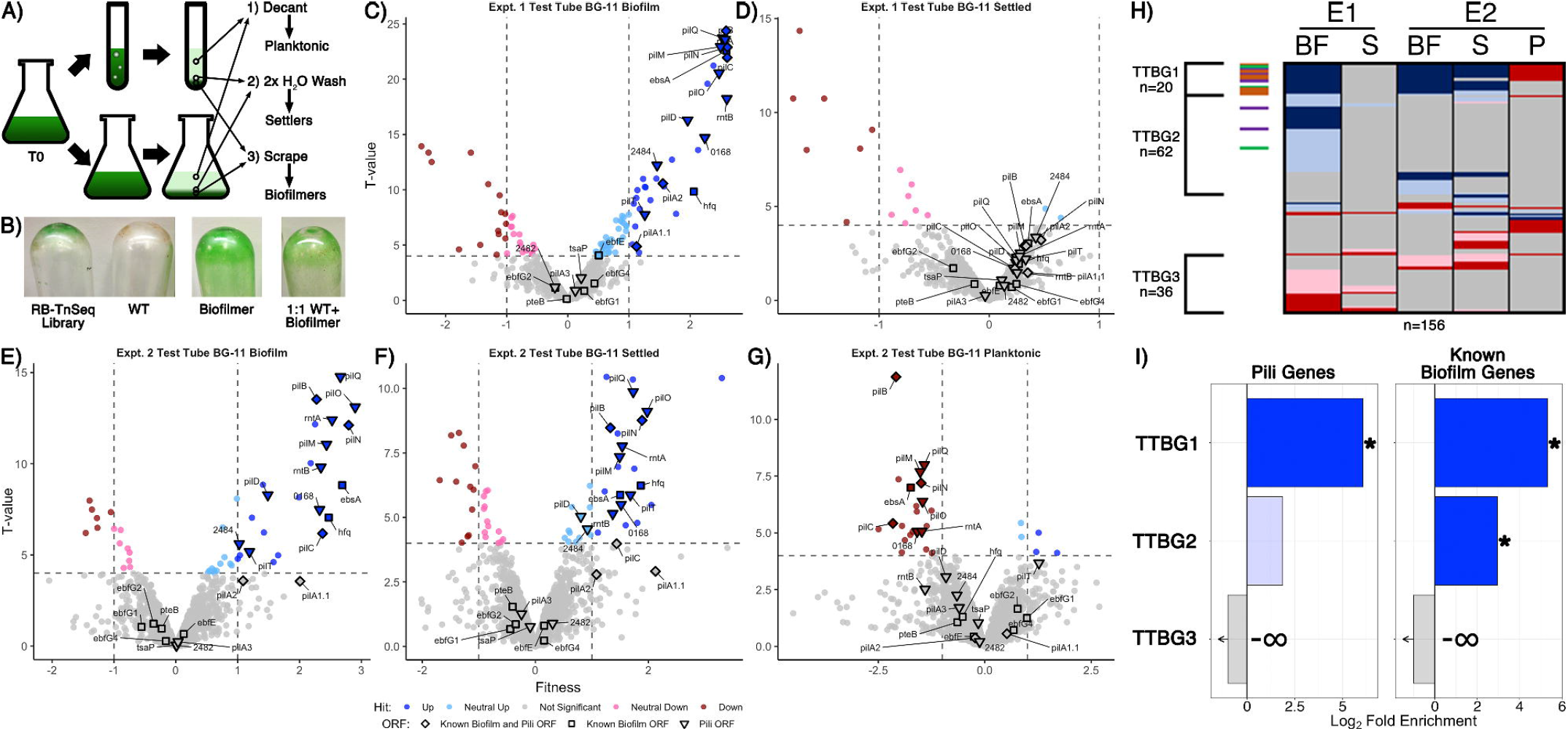
RB-TnSeq analysis of biofilming cultures of the pooled TnSeq library. A) Diagram of the RB-TnSeq library biofilm formation experiments and collection of samples. Triplicate samples were pooled prior to DNA extraction to ensure enough biomass and material for barcode sequencing. B) Representative biofilms, or lack thereof, left attached to the glass after decanting and washing steps for the RB-TnSeq library, WT, a homogeneous biofilmer culture, and a 1:1 mixture of WT and biofilmer. Volcano plots of experiment 1 (E1) test tube C) biofilm (BF) fractions and D) settled fractions (S), and experiment 2 (E2) test tube fractions in BG-11 E) biofilm, F) settler, and G) planktonic (P) fractions, presented as described for Figs. 1B,C except that the x-axis is the fitness value and the y-axis is the T-value statistic. H) Comparison of hit data for ORFs with significant fitness value changes, presented as described for Fig. 1D, for all BG-11 test tube fractions. I) Enrichment analyses for pili and known biofilm genes in the clusters of hits identified in H are presented as described for Fig. 1E.

Biofilmed library cultures were fractionated into three samples for barcode counting: (1) planktonic cells that were removed by decanting the media, (2) settled cells that were removed from the emptied test tubes with gentle water washes, and (3) biofilmed cells that required scraping to be removed from the glass culture vessels (Fig. 3A). The planktonic sample from one replicate experiment did not produce usable sequence data. For all other experiments, barcodes were counted and changes in mutant abundances were calculated based on statistical T-value thresholds^54^. The number of genes identified with statistically significant fitness value changes, called hits, in each fraction ranged from 16 to 103 (Fig. S1, File S4 Sheet 3), much smaller than the initial DEG ranges observed for RNA-Seq, but similar to the ranges of sDEGs acquired after filtering for strong fold changes (Figs. S1A,C). In total, 195 genes were classified as hits in at least one of the examined samples, with 108 having absolute fitness scores greater than 1. These hit counts are both substantially reduced compared to, and have little overlap with, the sDEGs in the RNA-Seq experiment (Fig. S10), indicating that the RB-TnSeq data set provides a distinctly different perspective on biofilm formation than does RNA-Seq.

Although the majority of genes, 1,725 or 89.8% of the 1,920 ORFs examined, show no significant fitness changes in any fraction examined (Figs. 3 & 4, File S4 Sheet 3), insertions in these genes were readily detected in all fractions (File S4 Sheet 1). These results indicate that the biofilm is composed not only of the mutants specially enriched in that fraction, but also of recruited or captured mutants that are also present in the planktonic fraction. It follows that a mutant that is under-represented in the biofilm may either be less fit to grow in the matrix or less likely to be recruited. Volcano plots demonstrate that more gene knockouts result in increased prevalence in the biofilm (positive hits) than decreased prevalence (negative hits) (Figs. 3C,E, & 4A,C,D). Positive hits in the fresh media test-tube experiments are dominated by known pili and biofilm genes whose loss enable film formation, such as *pilB*, while no pili or biofilm genes are present as negative hits in the biofilm fractions. Notably, the *ebfG* operon and *pteB* genes that prevent film formation when knocked out in a *PilB*::Tn*5* background show no significant changes in fitness in any of the experiments, indicating that these mutants can be incorporated into a biofilm even when they are not producing the critical elements for biofilm formation themselves (File S4 Sheet 5). Analysis of the settled fractions from both test tube experiments show inconsistent results (Figs. 3D, F). This inconsistency may be due to differences in the strength of the biofilms’ attachment to the glass vessel, so that wash steps differed in dislodging some of the biofilm as part of the settler sample. Nonetheless, the depletion of pili-related gene mutants and the *ebsA* mutant, which is known to lack pili^23^, from the planktonic fraction is consistent with the lack of pili resulting in faster rates of sedimentation^22^ (Fig. 3G, File S4 Sheet 5). Of the remaining 13 genes identified as strong negative hits (fitness < −1) in the planktonic fraction that are not known pili or biofilm associated genes (Table S6 group IG3, File S4 Sheet 8), 11 are predicted to encode proteins with secretion motifs (Table S6 group IG4) but were not detected previously in the exoproteomes of WT, *PilB*::Tn*5*, 1127Ω, or *PilB*::Tn*5* /1127Ω^22,27^. These genes either play some role in maintaining buoyancy or affect growth in the context of a biofilm.

**Figure 4:**
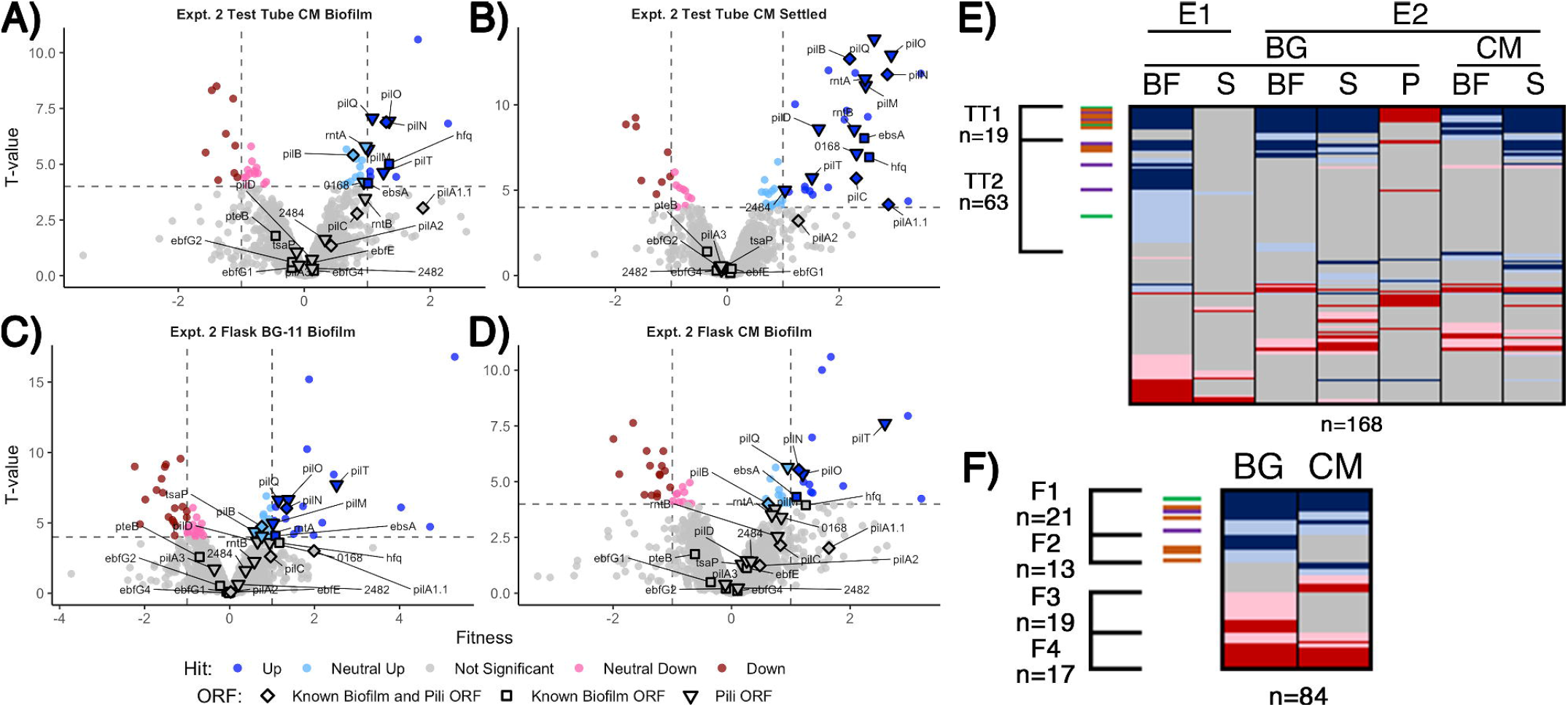
Conditioned media and flask RB-TnSeq analysis. Volcano plots of CM test tube A) biofilm and B) settled fractions, and biofilm fractions from flask experiments with C) BG-11 and D) CM, are presented as described for Figs. 1B,C. Comparison of fractions associated with E) all test tube experiments and F) flask experiments presented as described for Fig. 1D.

Comparative analysis of all fresh media test-tube experiments revealed three clusters of genes that represent consistent biofilm formers (TTBG1), and biofilm formers (TTBG2) or biofilm-depleted mutants (TTBG3) identified in one or more but not all of the experiments (Fig. 3H, File 4 Sheet 9). TTBG2 is enriched only in known biofilm genes (Figs. 3I & S8, File 4 Sheet 14), and TTBG3 lacks any known pili or biofilm genes. TTBG1 includes 6 new loci in addition to 14 known pili and biofilm mutants. Significantly, known biofilm genes that were never identified as sDEGs in the RNA-Seq analysis are present in TTBG1 (*pilD, hfq*, and *ebsA*) and TTBG2 (*ebfE* and *pilA2*). Thus, the remaining six genes in TTBG1, which include genes encoding a sigma factor, a response regulator, and a two-component sensor histidine kinase, represent likely candidates (Table S6 group IG5) for genes that participate in the self-suppression of biofilm formation, of which only Synpcc7942_0464, encoding a hypothetical protein, was a sDEG in the RNA-Seq analysis. The 59 novel hits in TTBG2 (Table S6 group IG6, File S4 Sheet 9) may also participate in the self-suppression of biofilm formation, prevent sedimentation, or prevent attachment to a growing biofilm because mutations that affect any of these processes would increase propensity for inclusion in the biofilm. TTBG2 is enriched for proteins conserved among cyanobacteria or involved in translation and ribosomal biogenesis (File S4 Sheet 14), and it is unclear how they affect biofilm formation. The 36 genes in TTBG3 (File S4 Sheet 9) whose mutants are absent from the biofilm are enriched for those encoding amino acid transport and metabolism, cell motility and chemotaxis, and protein maintenance genes (File S4 Sheet 14), consistent with phenotypes that may be needed for either recruitment into or survival in the biofilm.

In CM in test tubes, the settled fraction gave similar results to the biofilm fraction, where known biofilm and pili genes are enriched in the positive hits (Figs. 4A,B, & S8). Pattern analysis across all test tube experiments identified those that are unresponsive (TT1) and responsive (TT2) to CM (Fig. 4E, File S4 Sheet 13). The majority of the known biofilm and pili genes are found in TT1, including *pilB* whose homogenous mutant culture does not produce biofilms in CM^21^. We propose two hypotheses to explain the existence of transposon insertion mutants that are capable of producing biofilms, even in the presence of CM: 1) the self-suppressor generated by the WT-like cells in the transposon library experiments and in the mixed WT and mutant cultures was not at a high enough concentration to prevent biofilm formation at early time points, thus indicating a concentration-dependent window for enacting biofilm suppression, or 2) planktonic cells that would not initiate biofilm formation can be recruited into a mutant-seeded biofilm, which in CM would be recruitment of CM-responsive mutant cells into a biofilm seeded by CM-unresponsive mutants. Although both hypotheses may contribute to the reduced impact of CM on heterogeneous cultures in this experiment, the latter is supported here as this analysis highlights nine novel genes identified as media-independent biofilmers (Table S6 group IG7), a phenotype that was not previously encountered with *S. elongatus* biofilm mutants. This group includes a glycosyltransferase (Synpcc7942_0388) predicted to function in lipopolysaccharide production^73^. The identified genes are typified by the potential to impact the properties of the cell surface and thus impact biofilm formation or incorporation. Additionally, the analysis identified three consistent biofilmers that are responsive to CM, which include *sigF1*, Synpcc7942_0051, and Synpcc7942_B2646, and five mutants that biofilm only in CM, which includes a plasmid-encoded OmpR response regulator (Synpcc7942_B2647) (Table S6 groups IG 8 & IG9, File S4 Sheet 13).

Many known loci that were identified in test-tube assays did not display any significant fitness value changes in flasks (Figs. 4C,D, File S4 Sheets 5, 11, & 12). Subtle differences between the assays, including shearing stresses and increased gas exchange associated with bubbling, may alter division rates in the biofilm or the overall strength of the biofilm and affect fitness scores. Fresh BG-11 in flasks resulted in 16 novel biofilm formation mutants that did not form biofilms in BG-11 test tubes (Table S6 group IG10, File S4 Sheet 11), including *tsaP* and 3 mutants that had reduced prevalence in one of the test tube biofilms (Table S6 group IG11). Of these, five are genes involved in O-antigen production and susceptibility to grazing by amoeba, and were previously reported to result in cell aggregation, but not biofilm formation^34,74^ (Table S6 group IG12). We hypothesize that under the flask biofilming conditions, surface alterations that enhance aggregation or settling can more frequently enable incorporation into the heterogeneous biofilm.

Of the 13 flask biofilmers that were responsive to CM (group F2, Fig. 4F, File S4 Sheet 12), only *pilM, rntA*, and *tsaP* are known pili genes and nine are specific to flask biofilm formation (Table S6 group IG13). Eight mutants are specifically enriched in the flask biofilm in the presence of CM that were not otherwise enriched in test tube biofilms or in fresh BG-11 in flasks (Table S6 group IG14), including three co-transcribed chemotaxis-like genes of the *tax2* operon (Table S6 group IG15), whose function is unknown except that they are not involved in phototaxis in PCC 3055^25^.

### Validation of novel genes enabling or repressing biofilm formation

Together, the RNA-Seq and RB-TnSeq analyses identify hundreds of potential genes of interest. To evaluate the efficacy of these two approaches towards novel target identification, we selected a subset of previously untested genes to knock out in WT or in *PilB*::Tn*5* for biofilm evaluation, prioritizing those loci for which Unigene Set (UGS) transposon insertion vectors are available^30,31^. Of the 91 mutants attempted in WT, 79 produced segregated knockout mutants of the intended gene of interest (Table S7 & S8). Of these, we assayed candidates that were identified by: RNA-Seq only (30, 38%), RB-TnSeq only (38, 48%), or by both RNA-Seq and RB-TnSeq (11, 14%) (Table S7). Biological replicates of mutants were tested in bubbling test tubes, 96-well plates, and sometimes in flasks to increase the throughput of the biofilm assays and because different vessels impact the inclusion of a mutant in the biofilm. Most of the mutants created in a WT background did not form biofilms (49 of the 79, or 62%). Candidates based on RNA-Seq alone accounted for a higher false hit rate with 23 of the 30 assayed (77%) compared to 21 of the 38 assayed (55%) based on RB-TnSeq data alone.

The remaining 30 mutants formed biofilms to some degree and were classified based on the strength, reproducibility, and diversity of vessels in which biofilms were formed. 16 resulted in more replicates without any biofilms than with and were deemed “ mostly WT-like”. Of the remaining 14 mutants, most produced biofilms only sporadically or in a vessel-dependent manner. Only four mutants consistently produced strong, biofilms in multiple vessel assays – the predicted TPR-repeat lipoprotein with an O-linked N-acetylglucosamine transferase domain (Synpcc7942_0051) that was identified in the TTBG1 RB-TnSeq group; a gene predicted without experimental confirmation by Taton, et al.^24^ to encode the pilus assembly protein PilP (Synpcc7942_0168), whose impact on biofilm formation was not previously known; the gene encoding the RNA polymerase sigma factor SigF (Synpcc7942_1510); and the gene encoding the pilus assembly protein PilQ (Synpcc7942_2450). Notably, all genes identified from the TTBG1 RB-TnSeq group of interest whose validated mutations produced strong biofilm phenotypes were either never identified as sDEGs in the RNA-Seq analysis or had patterns of expression that were inconsistent with previous biofilm mutants (Tables S6, group IG16, & S7).

Similarly, 13 mutations were chosen for investigation in a *PilB*::Tn*5* background and assayed for mutants that no longer produce biofilms. Although none of the mutations completely abolished biofilm formation, knockout of a deacetylase (Synpcc7942_1393) (DEG in RNA-Seq group O-A4 full, Table S9, File S3 Sheet 15) or a pimeloyl-ACP methyl ester carboxylesterase (Synpcc7942_0774) decreased the reliability or strength of biofilm formation.

### Transcriptomics and phenomics reveal novel information on the biofilm state and the quality of the genome model

The RNA-Seq time course data and enrichment analysis results highlight phenotypic changes associated with adjusting to the self-shaded, nutrient and gas exchange-reduced biofilm lifestyle, including general stress responses, changes in iron, sulfur, and nutrient transportation and metabolism, protein maintenance, and energy generation through photosynthesis. The absence of mutants affected in many of these same genes in the biofilm fractions of the RB-TnSeq experiments support the hypothesis that these functions are essential for survival in biofilms. Given the substantial change in the cells’ niche as it forms biofilms, it is not surprising that a large portion of the transcriptome changes to properly adapt the cell for survival in the biofilm state, as has been observed in other systems^34,35,75^.

Strand-specific transcriptional data provided here and in previous publications^64,65^ can improve the genome models of PCC 7942 and related strains through resolving discrepancies between the observed transcriptome and the current genome annotation. The data confirmed that *pilB* is the first of four genes expressed as a single transcriptional unit (Fig. S9A)^65^. PilB::Tn5 transcript levels were reduced over 27 fold for the portion of *pilB* following the insertion site and the downstream genes including *pilT* and *pilC*, suggesting a polar effect. Phenotypic complementation, however, of PilB::Tn5 by introduction of *pilB in trans*^23^ indicates sufficient expression of the downstream genes in the mutant. Transcriptional coverage of the *hfq* gene does not exceed one read count until positions annotated as codons 25 or 27, depending upon the strain analyzed, suggesting that an alternative GTG start codon previously designated as amino acid position 28 begins the open reading frame (Fig. S9B). This shorter annotation is consistent with the NCBI conserved domain database’ identification of an Sm-like RNA binding domain from codons 35 to 95^76^.

### RNA-Seq compared with RB-TnSeq

As the results for *hfq* and *ebsA* highlight, some key genes may not be identified by transcriptomic approaches because protein levels in cyanobacteria are poorly correlated with transcript levels^77^ or protein functions are regulated post-transcriptionally. In contrast, RB-TnSeq directly probes the connection between genotype and phenotype in a high-throughput manner^54,78^. In this project, we leveraged the self-fractionation of the sample that occurs through biofilm formation as an application for an RB-TnSeq library. In comparison to experiments where changes in fitness score reflect mutants’ rate of division relative to the rest of the library’s population in the presence of a selective condition^24,53,54,72,78^, this application’s fitness scores are also affected by the capacity to reproducibly fractionate the cultures as well as mutation- and niche-dependent variations in rates of growth, survival, recruitment of WT-like cells into the biofilm, and settling, which is more reflective of the ability to remain planktonic rather than the capacity to generate biofilm structures.

Nonetheless, the RB-TnSeq strategy provides a more direct identification of both known and novel gene products that participate in a complex behavior such as biofilm formation, even with fewer experimental samples, than does RNA-Seq. This conclusion is based on the higher rate of validation of hits for the RB-TnSeq data than the RNA-Seq data and the lack of known biofilm regulatory genes in the transcriptomics data (Figs. S1 & S10). Because RB-TnSeq experiments require less sequence coverage than RNA-Seq samples, this strategy is also a less expensive methodology for identifying novel candidates involved in nuanced and complex behavior.

Because the WT *S. elongatus* PCC 7942 background constitutively represses biofilm formation, the experimental design would not identify mutants that are defective in components that are required for biofilm formation, such as the *ebfG*-*pteB* genes. These genes show the highest fold changes in transcript expression related to biofilm formation, but their inactivation in the RB-TnSeq library showed no significant changes in fitness values, likely due to the secretion of their encoded proteins by other biofilm forming mutants in the library. Future applications of Interaction RB-TnSeq (IRB-Seq), in which a second mutation (e.g., *pilB*::Tn*5*) is introduced into the RB-TnSeq library^72^ to stimulate biofilm formation, or an RB-TnSeq library constructed in a native biofilmer such as UTEX 3055, is more likely to identify genes that are involved in biofilm formation process rather than the self-suppression mechanism.

### Novel genes of biofilm regulation in *S. elongatus*

Of the large number of potential candidates for participating in regulation or formation of biofilms, only four resulted in strong and consistent biofilm phenotypes: Synpcc7942_0051, Synpcc7942_0168, *sigF* (Synpcc7942_1510), and *pilQ* (Synpcc7942_2450). Synpcc7942_0051 shares a domain with proteins in heterotrophic bacteria that are transcriptionally regulated by a sigma-54-interacting response regulator and a histidine kinase and participate in exopolysaccharide production and protein export in biofilm forming bacteria^79^. If this protein plays a similar role in PCC 7942 biofilms, then histidine kinases and response regulators that were identified as potential genes of interest (Table S6 group IG17) are candidates for participating in the regulatory circuit for controlling biofilm self-suppression. These validated hits, combined with the large set of genes with unknown function highlighted by the transcriptome and phenome data sets, serve as a starting point for future investigations of novel mechanisms of biofilm regulation in *S. elongatus* PCC 7942.

## Acknowledgments

Studies were supported by the program of the National Science Foundation and the US-Israel Binational Science Foundation (NSF-BSF 2012823 to R. Schwarz and S. Golden). This work was also supported by grants from the Israel Science Foundation (ISF 1406/14 and 2494/19) to Rakefet Schwarz.

## Legends to Supplemental Material

**Figure S1. Comparing the results of RNA-Seq and RB-TnSeq analyses**. Combined scatter and box plots of A) the number of DEGs for the pairwise comparisons presented in Fig. S1 or the number of hits from all RB-TnSeq fractions, B) the percent of genes investigated in each experiment that are DEGs or RB-TnSeq hits, C) the number of sDEGs or RB-TnSeq hits, and D) the percent of genes investigated in each experiment that are sDEGs or RB-TnSeq hits. Boxed letters indicate groups that are not significantly different (p > 0.05) based on pairwise two-tailed t-tests assuming unequal variances; RB-TnSeq results are significantly different from those of RNA-Seq when considering all DEGs, but not when limiting the analysis to sDEGs.

**Figure S2. Volcano plots for all ten pairwise comparisons relevant to the tested RNA-Seq experimental variables**. Plots are presented as described in Figs. 1B,C and are derived from either the A) DESeq analysis or B) Rockhopper analysis of RNA-Seq data. For the Rockhopper analysis, the software returns FDR values of 0.0 for a number of ORFs, which become inifinite values on the y-axis’ log-scale. These infinite data points are accumulated at the top of the graph.

**Figure S3. Enrichment analyses of A) pili genes and B) known biofilm genes in all named RNA-seq clusters of interest**, as identified in Figs. 1 and 2 and File S3. The graphs are presented as described for Fig. 1E.

**Figure S4. Enrichment analyses of sDEGs groups**, as identified in Fig. 1D. The graphs are presented as described for Fig. 1E, except only categories of information accumulated in Table S3 with at least one gene present in the interest group are shown.

**Figure S5: Day 4 and time course RNA-Seq enrichment analyses**. Functional enrichment analysis of Cyanobase COGs for A) Day 4 and B) over time groups of interest as identified in Fig. 2D and E. Colors identify the data for the appropriate group of interest, as provided in the graph legend.

**Figure S6. Enrichment analyses of all Day 4 sDEGs groups**, as identified in Fig. 2D. The graphs are presented as described for Fig. 1E, except only categories of information accumulated in Table S3 with at least one gene present in the interest group are shown.

**Figure S7. Enrichment analyses of all over time sDEGs groups**, as identified in Fig. 2E. The graphs are presented as described for Fig. 1E, except only categories of information accumulated in Table S3 with at least one gene present in the interest group are shown.

**Figure S8. Enrichment analyses** of A) pili genes and B) known biofilm genes in all named RB-Tnseq clusters of interests, as identified in Figs. 3 and 4 and File S4. The graphs are presented as described for Fig. 1E.

**Figure S9: RNA-Seq Coverage of *pilB* operon and *hfq***. A) Strand-specific transcriptional coverages in the region of the *PilB*::Tn*5* insertion and the downstream operon in WT and *PilB*::Tn*5* samples in fresh BG-11 on Day 1. Both coverage tracks are at the same scale, with the highest peak being approximately 1000x. B) Strand-specific transcriptional coverages in the region of *hfq* (Synpcc7942_1926) in WT and *PilB*::Tn*5* samples in fresh BG-11 on Day 1. Both coverage tracks are at the same scale, with the highest peak being approximately 70x. Transcriptional coverage starts in the middle of the canonical annotation (black arrow) but before the proposed ORF based on an alternative start codon (black brackets) based on the location of a predicted Sm-like RNA binding domain (purple).

**Figure S10. Venn Diagrams of Hit Sets for RNA-Seq and RB-TnSeq data sets**. A) Strong DEGs (sDEGs) for RNA-Seq versus RB-TnSeq Hits, B) sDEGs for RNA-Seq versus RB-TnSeq strong hits, C) DEGs for RNA-Seq versus RB-TnSeq hits, and D) DEGs for RNA-Seq versus RB-TnSeq strong hits. Areas are proportional to numbers provided for overlaps, RNA-Seq only DEGs, or RB-TnSeq hits.

**Supplemental Table S1. List of Genes Known to Impact Biofilm Formation**

**Supplemental Table S2. List of Pilin Genes from Taton et al. (2020)**

**Supplemental Table S3. Functional Categorization, Essentiality, Conservation, and Signal Peptide Prediction for ORFs in Synechococcus elongatus PCC 7942**

**Supplemental Table S4. Rockhopper Predicted Novel RNAs**

**Supplemental Table S5. Summary of Hits of Interest**

**Supplemental Table S6. Groups of Genes Discussed in the Text Not Listed in a Named Interest Group**

**Supplemental Table S7. Biofilm Assay Results for Single Mutants in a WT Background**

**Supplemental Table S8. Mutations attempted in WT that did not produce segregated clones**

**Supplemental Table S9. Biofilm Assay Results for *PilB*::Tn*5* Double Mutants**

**Supplemental Table S10. Vectors and Associated Primer Pairs Used in This Study**

**Supplemental File 1. Rockhopper RNA-Seq Analysis**. Sheets include 1) alignment statistics, 2) all data output by the Rockhopper software, 3) raw counts, 4) normalized counts, 5) RPKM and expression values, 6) fold and statistical values for all pairwise comparisons of samples, 7) a summary of all DEG assignments, 8) lists of all DEGs in each pairwise comparison, 9) a summary of DEG assignments for predicted RNAs, and 10) a summary of DEG assignments for known biofilm and pili genes,

**Supplemental File 2. DESeq RNA-Seq Analysis**. Sheets include 1) alignment statistics,

2) raw counts, 3) normalized counts, 4) ANOVA analysis for DEGs across all samples, 5) fold and statistical values for all pairwise comparisons of samples, 6) a summary of all DEG assignments, 7) lists of all DEGs in each pairwise comparison, and 8) a summary of DEG assignments for known biofilm and pili genes,

**Supplemental File 3. RNA-Seq Analysis**. Sheets include 1) Consistency analysis or the data for generating the union of all pairwise comparisons from Rockhopper and DESeq data, 2) a summary of all DEG assignments, 3) lists of all DEGs in each pairwise comparison, 4) a summary of sDEGs for the 10 relevant pairwise comparisons, 5) lists of sDEGs in each of the 10 relevant pairwise comparisons, 6) a summary of DEG assignments for known biofilm and pili genes, 7) comparison data of DEGs for Day 1 pairwise comparisons, 8) comparison data of sDEGs for Day 1 pairwise comparisons used to produce the middle graph of Fig. 1D, 9) comparison data of DEGs for Day 4 pairwise comparisons, 10) comparison data of sDEGs for Day 4 pairwise comparisons used to produce Fig. 2D, 11) comparison data of DEGs for Day 4 versus Day 1 pairwise comparisons, 12) comparison data of sDEGs for Day 4 versus Day 1 pairwise comparisons used to produce Fig. 2E, 13) comparison data of DEGs for WT BG-11 Day 4 versus Day 1 against *PilB*::Tn*5* BG-11 Day 1 vs WT BG-11 Day1, 14) comparison data of sDEGs for WT BG-11 Day 4 versus Day 1 against *PilB*::Tn*5* BG-11 Day 1 vs WT BG-11 Day1 used to produce the left graph of Fig. 1D, 15) the reduced set of DEGs for Day 1 comparison data, 16) the reduced set of sDEGs for Day 1 comparison data used to produce the right graph of Fig. 1D, 17) enrichment analysis data for named DEG clusters of interest for all gene categories listed in Table S3, and 18) enrichment analysis data for named sDEG clusters of interest for all gene categories listed in Table S3.

**Supplemental File 4. RB-TnSeq Analysis**. Sheets include 1) Gene insertion counts, 2) Fitness and T-values, 3) a summary of all hit assignments, 4) lists of all hits in each sample, 5) a summary of hit assignments for known biofilm and pili genes, 6) a listing of genes as per a traditional venn diagram comparing various sample results, 7) comparison of experiment 1 samples, 8) comparison of experiment 2 BG-11 test tube samples, 9) comparison all test tube BG-11 samples, data used to produce Fig. 3H, 10) comparison of all experiment 2 test tube samples, 11) comparison of all BG-11 biofilm samples, 12) comparison of experiment 2 flask samples, data used to produce Fig. 4F, 13) comparison of all test tube samples, data used to produce Fig. 4E, and 14) enrichment analysis data for named hit clusters of interest for all gene categories listed in Table S3.

