## Supplementary figures and images for "Transcriptomic and phenomic investigations reveal elements in biofilm repression and formation in the cyanobacterium *Synechococcus elongatus* PCC 7942"

### Supplemental Figure 1

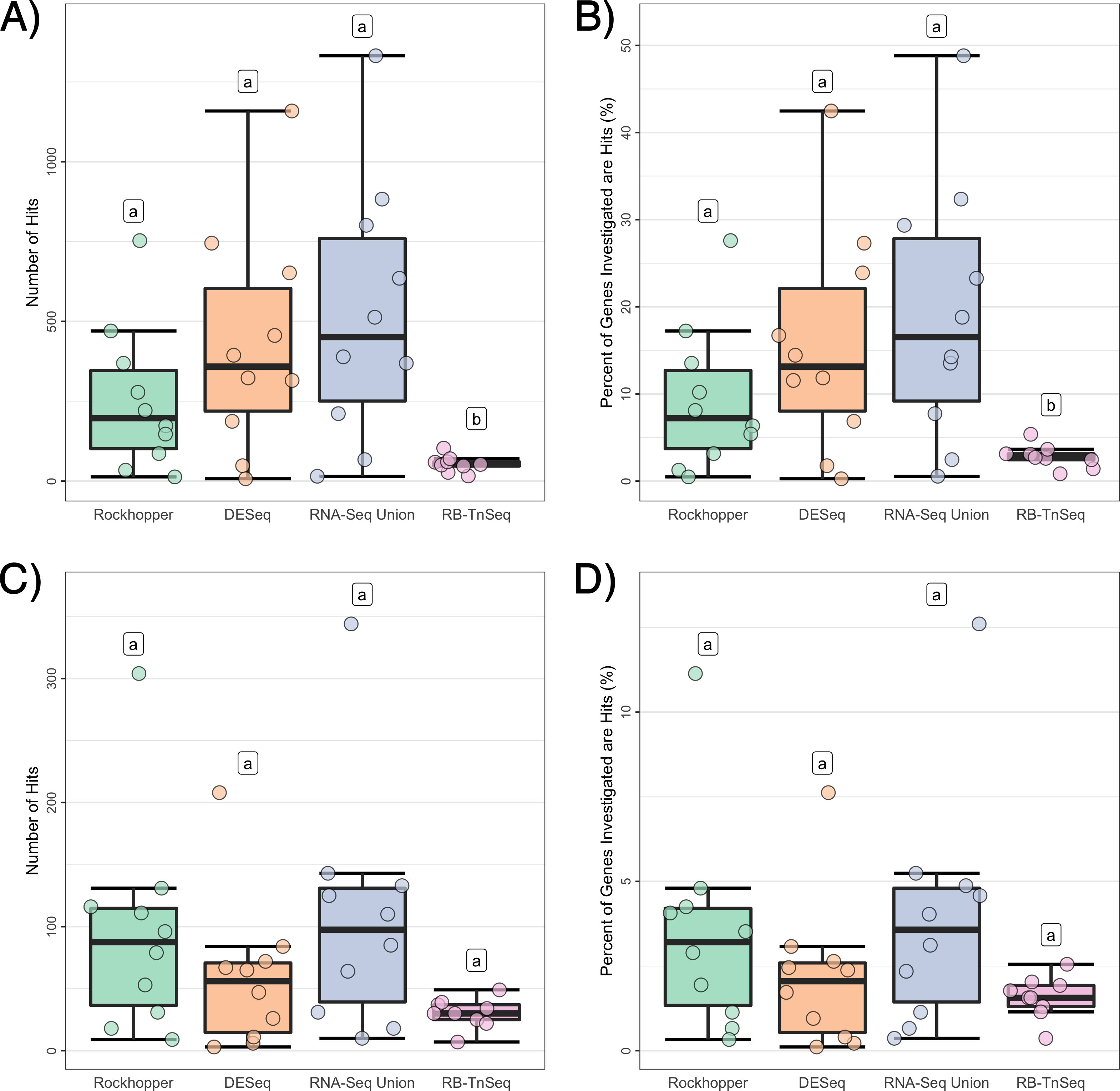

### Supplemental Figure 2

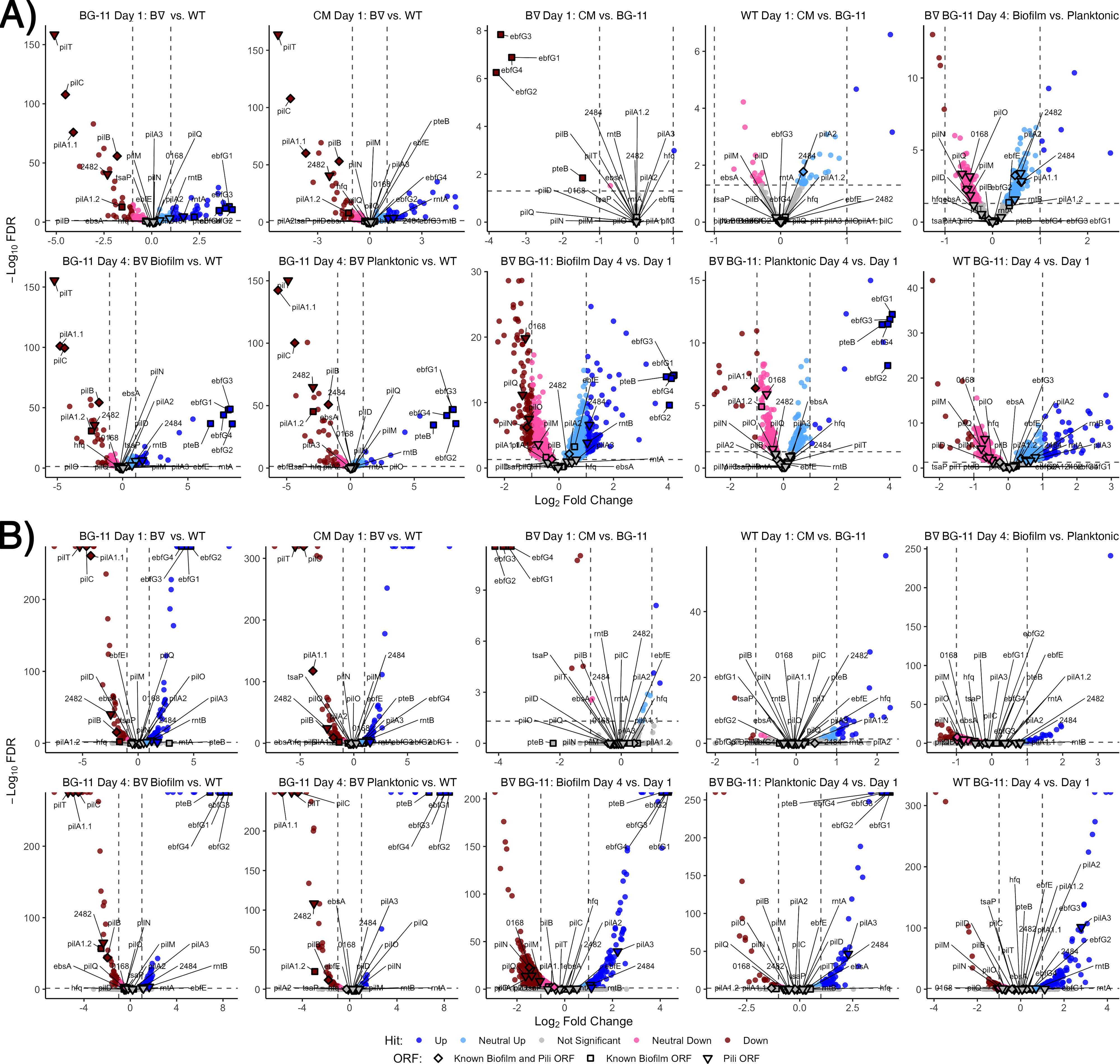

### Supplemental Figure 3

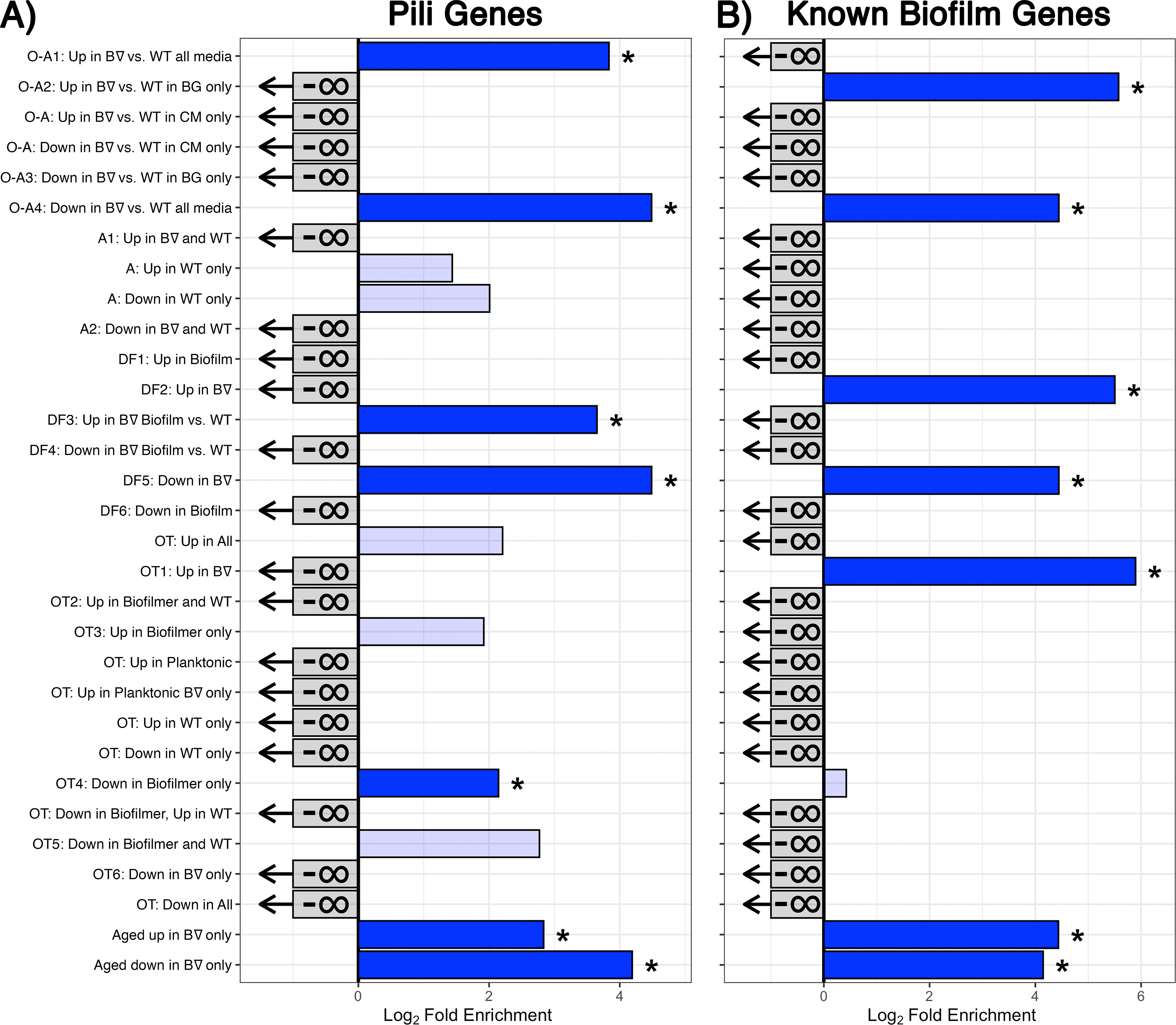

### Supplemental Figure 4

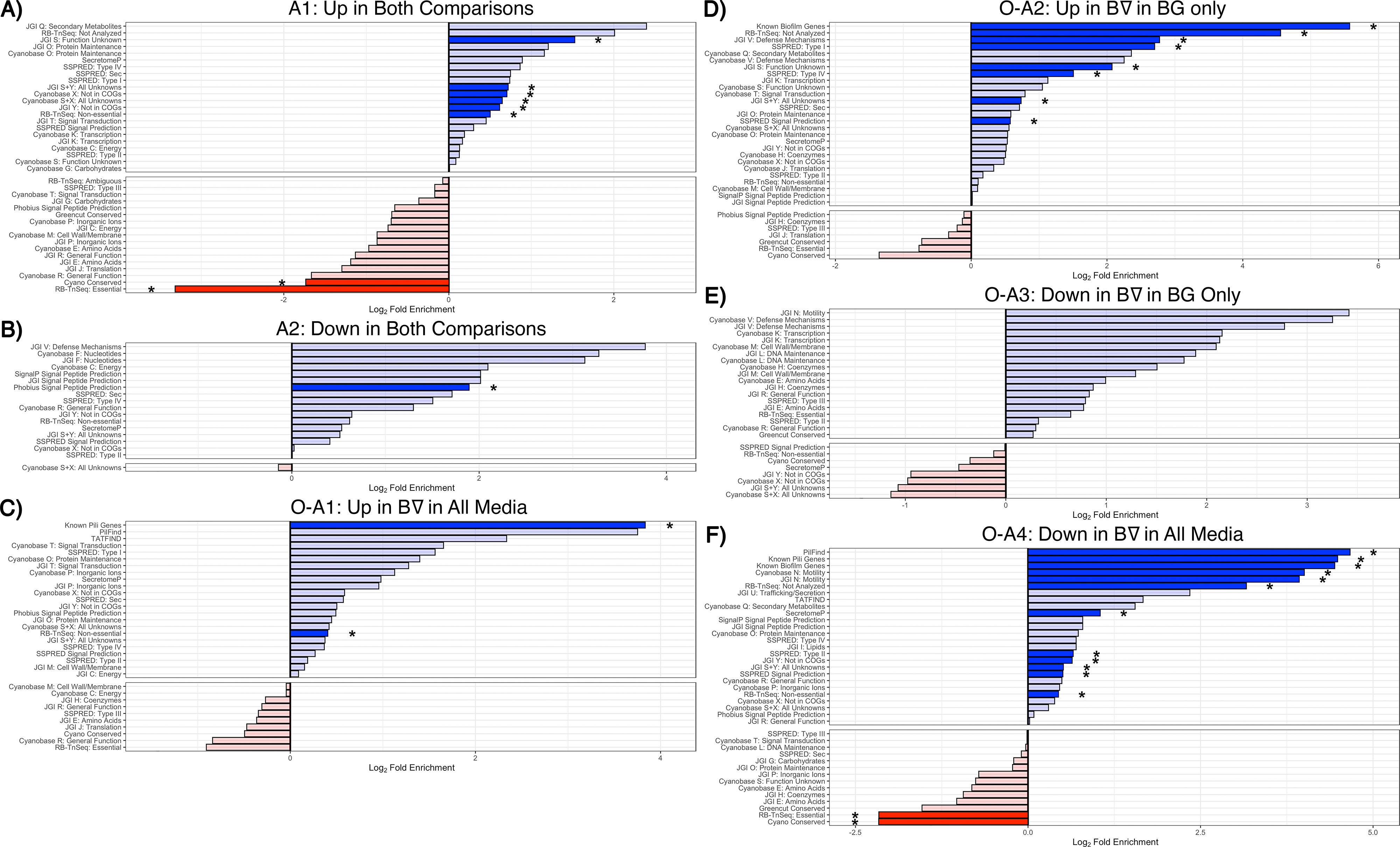

### Supplemental Figure 5

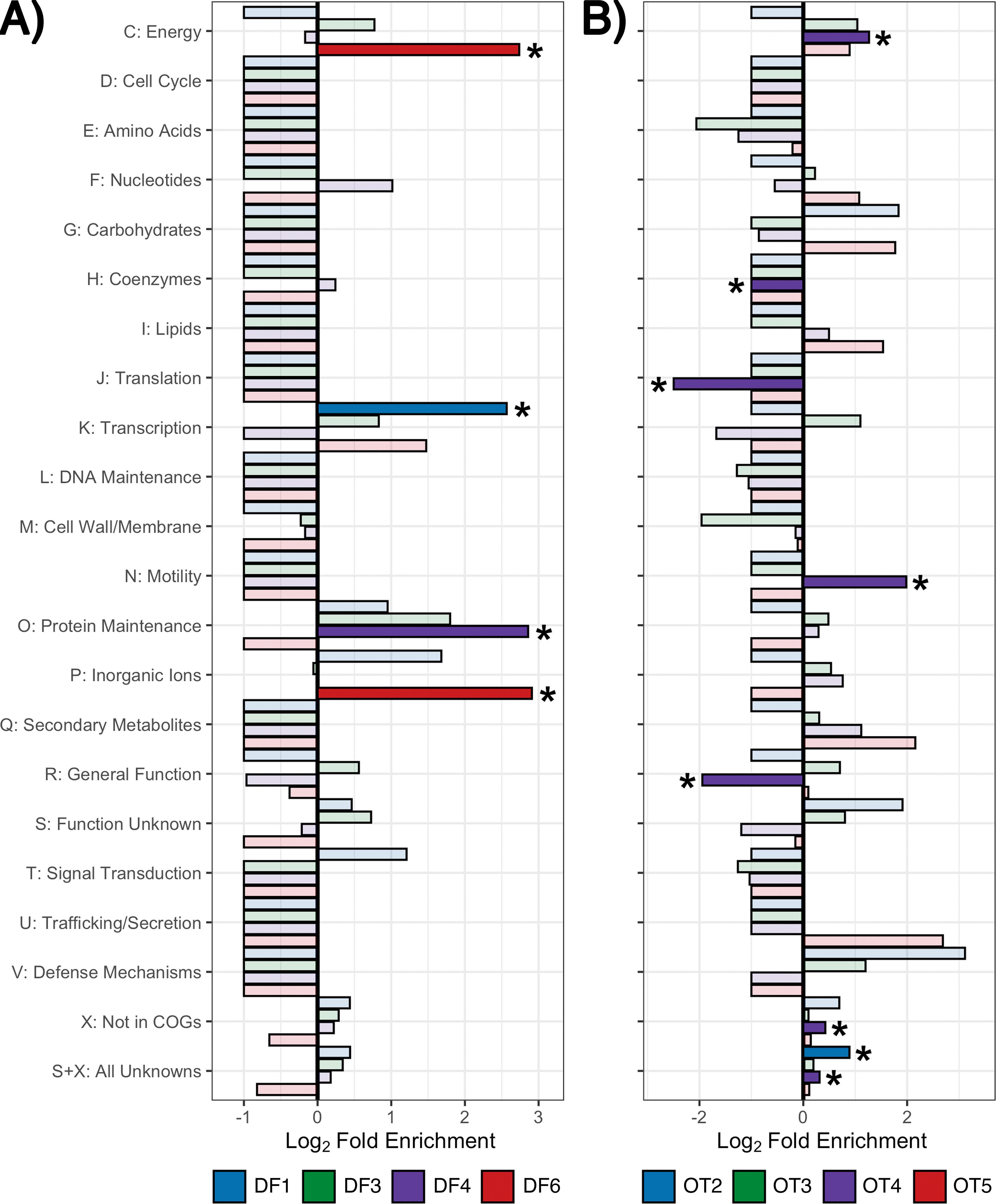

### Supplemental Figure 6

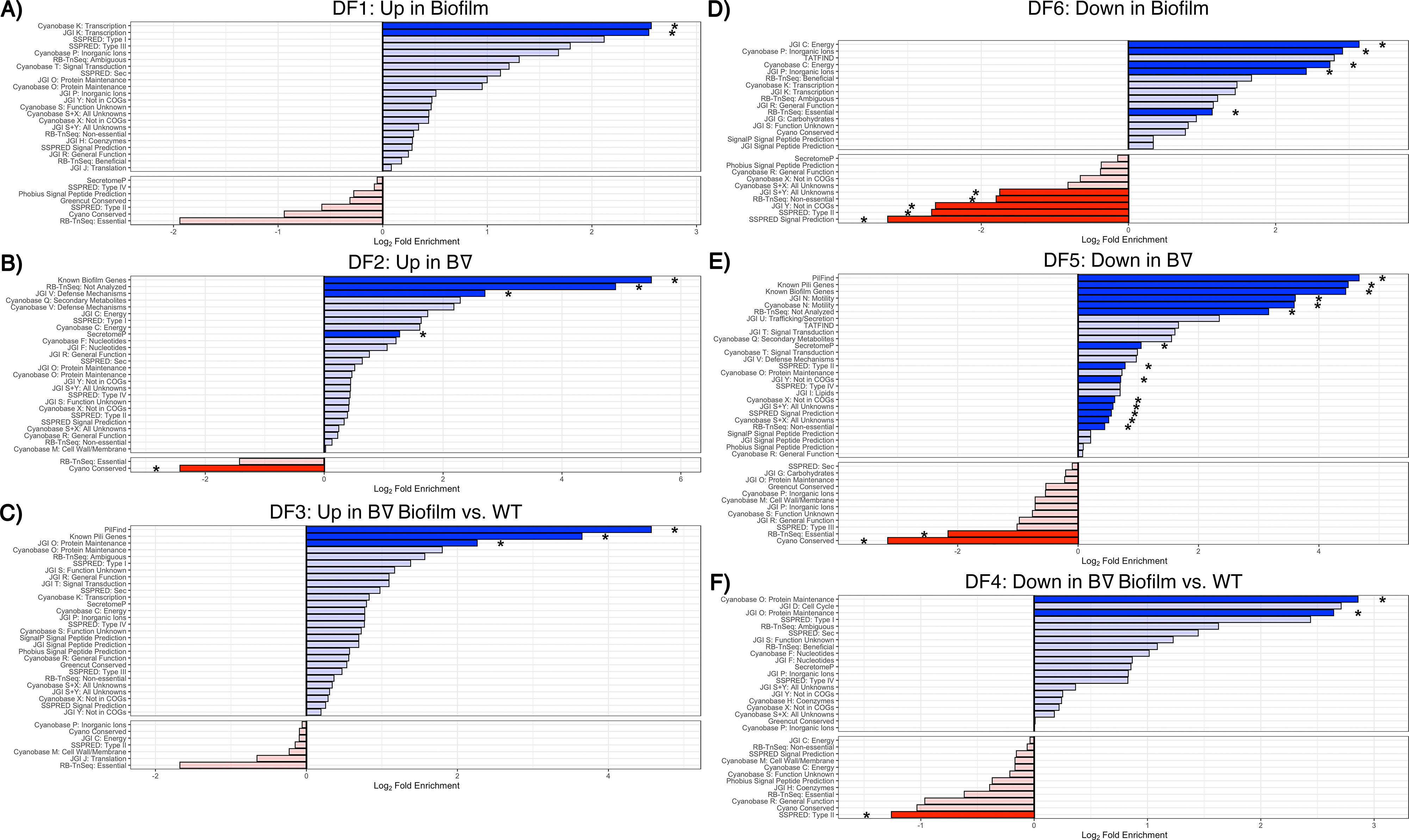

### Supplemental Figure 7

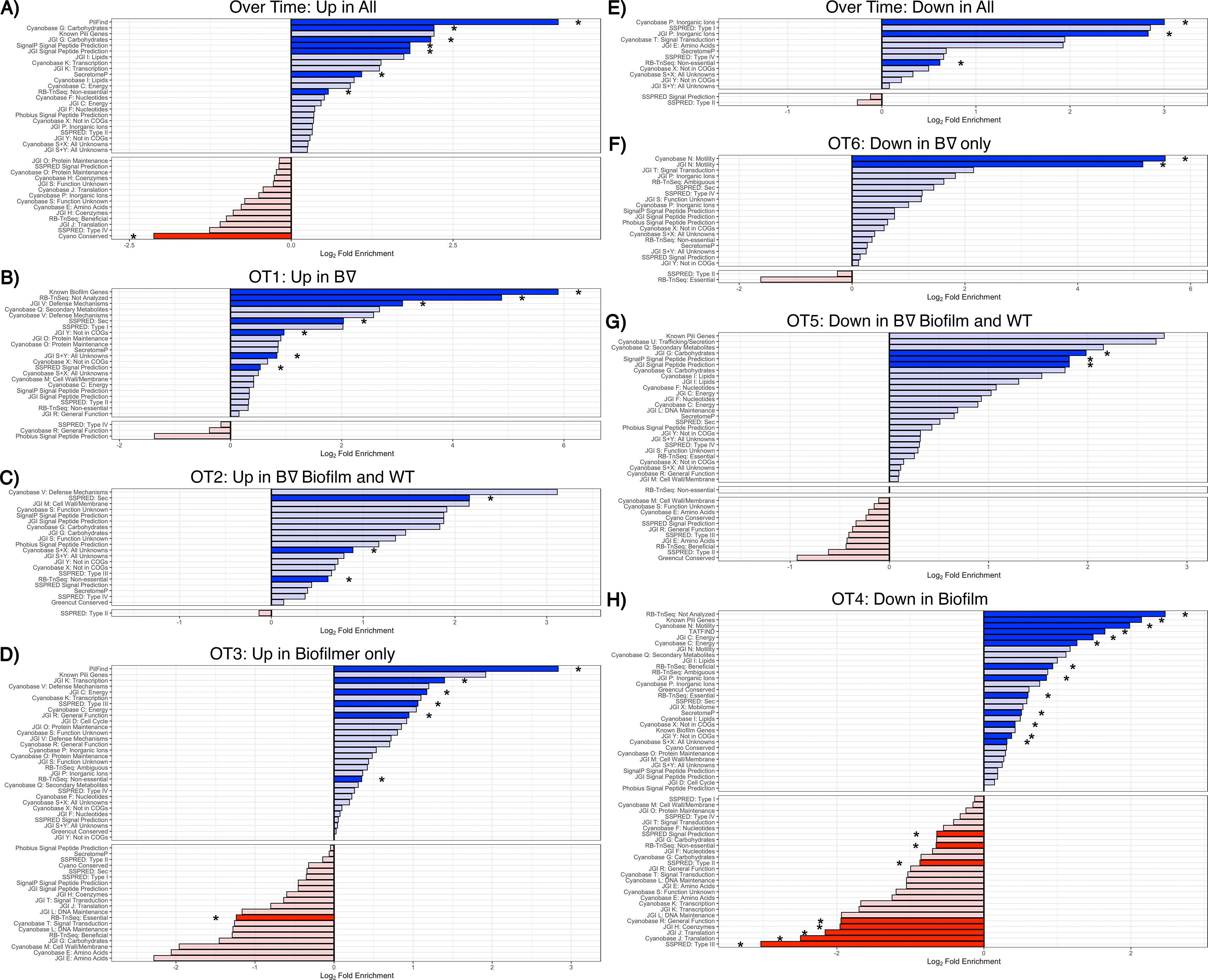

### Supplemental Figure 8

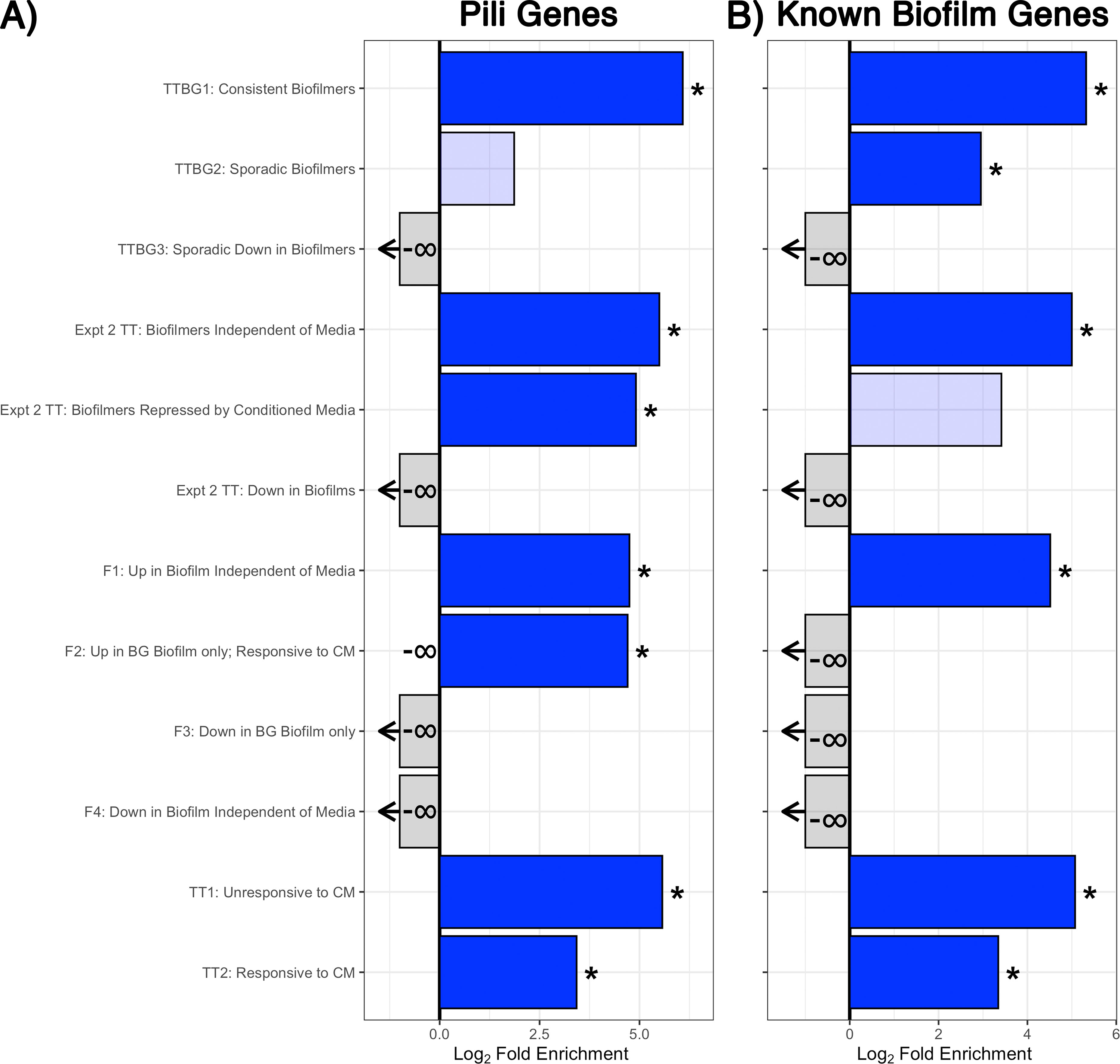

### Supplemental Figure 9

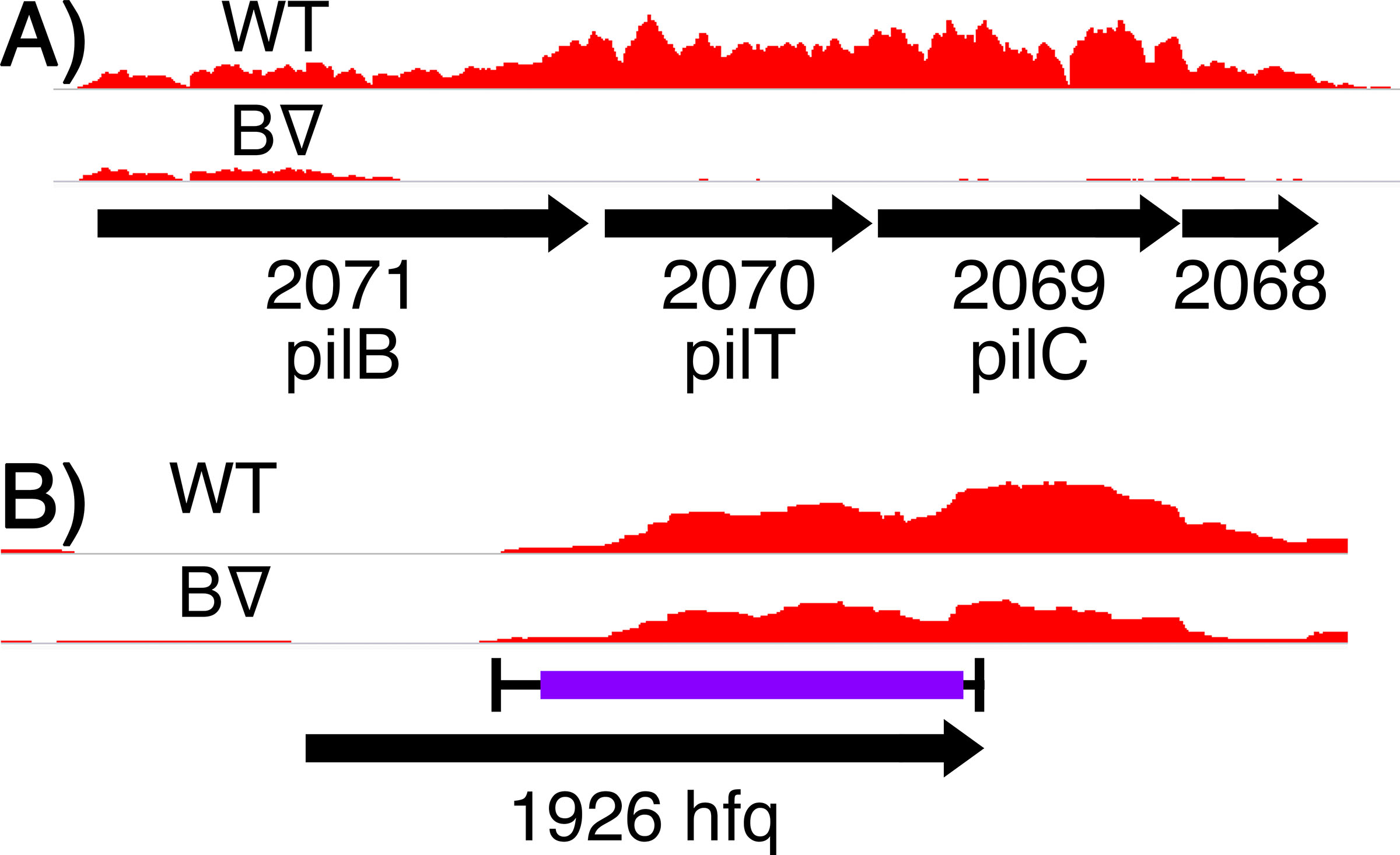

### Supplemental Figure 10

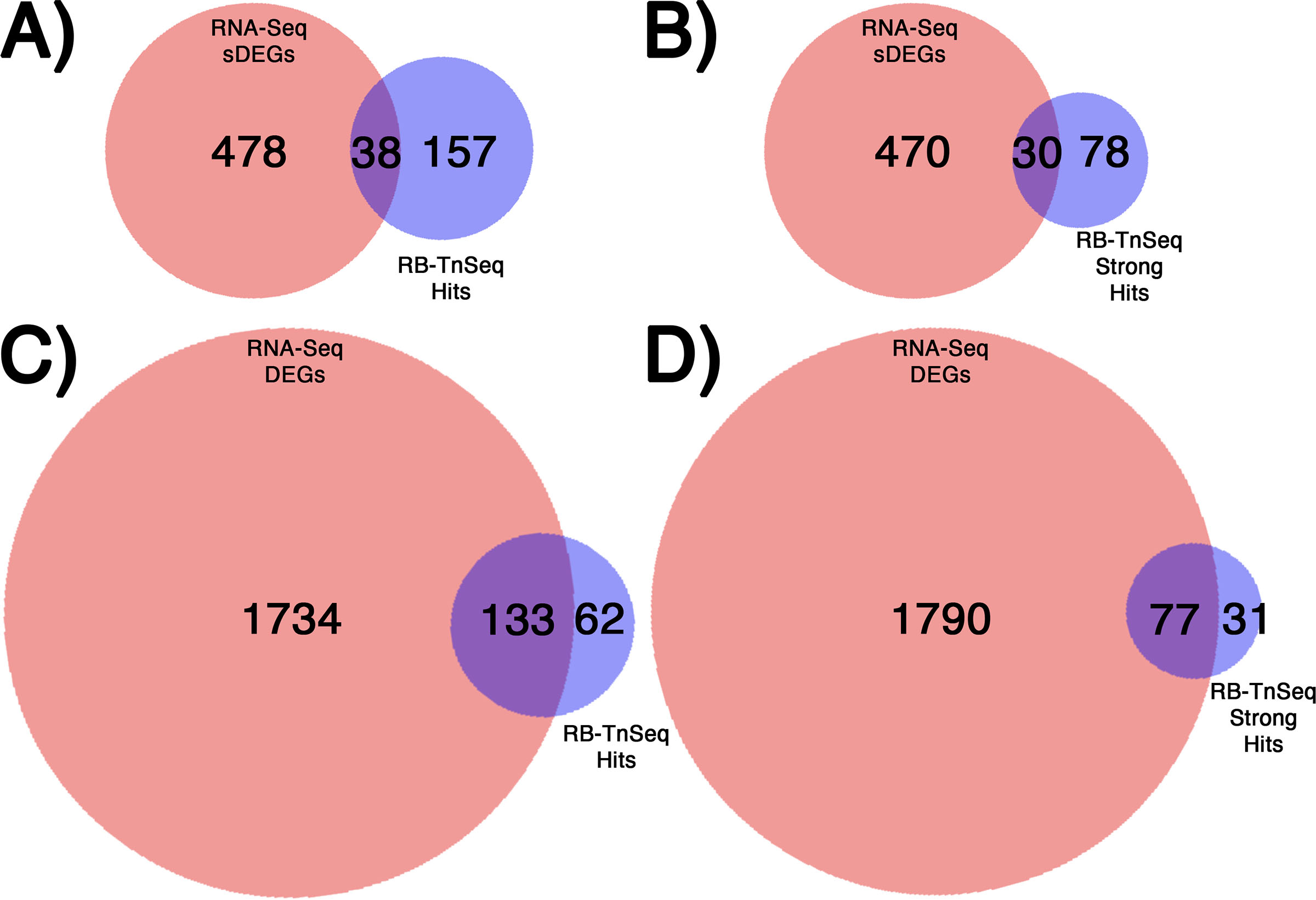
